# The cut site specificity of the influenza A virus endoribonuclease PA-X allows it to discriminate between host and viral mRNAs

**DOI:** 10.1101/2022.07.08.499385

**Authors:** Lea Gaucherand, Amrita Iyer, Isabel Gilabert, Chris H. Rycroft, Marta M. Gaglia

## Abstract

Widespread shutoff of host gene expression through RNA degradation is an advantageous way for many viruses to block antiviral responses. However, viruses still need to maintain expression of their own genes and host genes necessary for replication. The influenza A virus host shutoff endoribonuclease PA-X solves this problem by sparing viral mRNAs and some host RNAs. To understand how PA-X distinguishes between RNAs, we characterized PA-X cut sites transcriptome-wide. This analysis shows that PA-Xs from multiple influenza strains cleave RNAs at GCUG tetramers in hairpin loops. Importantly, GCUG tetramers are enriched in the human but not the influenza transcriptome. Moreover, optimal PA-X cut sites inserted in the influenza A virus genome are quickly selected against during viral replication. This finding suggests that PA-X evolved these cleavage characteristics to target host but not viral mRNAs, in a manner reminiscent of cellular self vs. non-self discrimination.

## Introduction

Many viruses block host gene expression to take over the infected cell. This process, termed host shutoff, is thought to promote viral replication by preventing antiviral responses and redirecting cellular resources to viral processes. Several viruses from divergent families accomplish host shutoff through RNA degradation by endoribonucleases (endoRNases), including the human pathogen influenza A virus, a negative-sense single-stranded RNA virus with a segmented genome^1–3^. This convergent evolution highlights the benefit of using endoRNases to induce host shutoff. In influenza A virus, the virus-encoded endoRNase PA-X drives widespread host RNA depletion^3, 4^. PA-X activity has critical consequences for the host, as infection with a PA-X-deficient virus induces more inflammation in animal models, often causing higher morbidity and mortality than wild-type infections^3, 5–8^. While immune modulation by PA-X *in vivo* has been widely documented, few studies have investigated its molecular mechanism of action. Moreover, the RNA targeting specificity of PA-X and its contribution to PA-X function during infection are still incompletely understood.

PA-X is produced from segment 3 of influenza A virus, which also encodes the polymerase acidic (PA) subunit of the influenza RNA-dependent RNA-polymerase (FluPol). PA-X is produced when a +1 ribosomal frameshift occurs after addition of amino acid 191^3, 9^. PA-X thus shares its N-terminal domain with PA, which has a PD-D/E-XK nuclease fold and endoRNase activity^10, 11^. PA uses this domain to “snatch” capped 5’ ends of host mRNAs to initiate viral mRNA transcription^10^, whereas PA-X uses it for host shutoff^4^, and *in vitro* studies suggest that the two proteins have different substrate preference^12^. We previously found that PA-X activity is not indiscriminate, as it down-regulates host RNAs synthesized by RNA Polymerase II (RNAPII) but not RNAPI and III^13^. Interestingly, this feature is shared by all characterized viral endoRNases and degradation factors that drive host shutoff^1, 14^. In influenza A virus, this specificity may be explained by a connection between PA-X and cellular RNA splicing, which may govern how PA-X reaches its target RNAs^4^. However, unspliced reporter RNAs can also be degraded to some extent, suggesting that there may also be other determinants. Additionally, unlike some other viral endoRNases, PA-X does not appear to consistently down-regulate viral mRNAs^13^, suggesting it can distinguish between host and viral transcripts. Viral RNA levels are unaffected by PA-X in cells infected with the influenza A/PuertoRico/8/1934 (H1N1) virus strain^13^, although some change was reported in the context of infections with the A/California/04/2009 (H1N1) strain^15^. Also, splicing alone does not explain the virus-host discrimination, as the mRNAs from the influenza A virus M and NS segments are spliced by the host machinery but spared by PA-X. These observations indicate that there are additional unknown components to PA-X selectivity.

While the role of cofactors in RNase specificity is well-established, cut site specificity remains poorly understood for many endoRNases, although sequence and/or structure preferences have been identified when it has been examined. For example, phage T4 RegB cleaves mRNA at GGAG sequences located in specific structures^16, 17^, and fungal α–sarcin cleaves 28S ribosomal RNA at GAGA sequences within loops^18^. However, the complexity of cut site motifs hampers their identification using classical *in vitro* approaches. Next generation sequencing has recently been used to tackle this challenge by profiling endoRNase cleavage products in cells, and for example, identify the complex preferred motif of *Salmonella* endoRNase E^19^. We have employed it to identify the cut site specificity of the Kaposi’s sarcoma-associated herpes virus (KSHV) endoRNase SOX^20^. Knowing the cut site specificity is critical to understand which RNAs are directly targeted by RNases and how.

While we have used RNA steady state levels to report on PA-X activity, this approach may be confounded by feedback effects on transcription^21^ and the inherent instability of certain RNAs. Thus, to better understand PA-X targeting, we directly probed PA-X cut sites throughout the transcriptome using 5’ Rapid Amplification of cDNA Ends (5’ RACE) adapted to high-throughput sequencing and a custom pipeline (PyDegradome)^20^. We report that PA-X preferentially cleaves host RNAs at GCUG sequences within hairpin loops, likely after splicing. Importantly, these preferred cleavage motifs are more abundant in host than viral mRNAs. Moreover, inserting a preferred PA-X cleavage sequence in a viral segment is detrimental to viral growth in the presence, but not in the absence, of PA-X. These findings suggest that PA-X cut site specificity contributes to its ability to broadly target the host transcriptome while sparing the virus, and serves as a “self/non-self” discrimination mechanism for influenza A virus.

## Results

### Identification of PA-X cut sites transcriptome-wide

In preliminary 5’ RACE experiments, we found that PA-X cuts reporter mRNAs at a specific location (data not shown). We thus hypothesized that PA-X cleaves host RNAs at distinct places in the transcriptome, contrary to what we previously thought^13^. To test this hypothesis transcriptome-wide, we used 5’ RACE coupled to high-throughput sequencing (5’ RACE-seq, **Fig. 1A**) and compared RNA fragments from mock infected cells to cells infected with the wild-type (WT) influenza A/PuertoRico/8/1934 (H1N1) virus strain (henceforth “PR8”), or PR8 engineered to lack PA-X (PR8-PA(ΔX) or ΔX, **Extended Data Fig. 1A**^4^). The mutations in PR8-PA(ΔX) introduce a stop codon in the PA-X +1 reading frame and should reduce the amount of frameshifting, abolishing production of full-length PA-X^4^. They are silent in the PA 0 reading frame, and do not affect PA protein levels (**Extended Data Fig. 1B**). While we lack a reliable antibody to measure PR8 PA-X, we have previously profiled the transcriptional changes due to these mutations and confirmed that PR8-PA(ΔX) causes reduced host shutoff compared to WT PR8^4^. Since the host exonuclease Xrn1 degrades RNA fragments following cleavage by PA-X^13^, we used Xrn1 knock-out human lung A549 cells to enrich for PA-X fragments^22^ (**Extended Data Fig. 1C**). We confirmed by qRT-PCR that the samples expressed similar levels of viral genes, and that host shutoff was detected in the WT PR8 infected samples using the PA-X target G6PD mRNA as a readout^13^ in WT and Xrn1 knock-out A549 cells (**Fig. 1B-C**). For 5’ RACE-seq, we ligated an RNA adapter to the 5’ phosphate at the end of cleaved RNAs, which include PA-X cleaved RNAs, and prepared sequencing libraries (**Fig. 1A**). After aligning reads to the human genome, the junction between the 5’ RACE adapter and the human sequence represents the position where PA-X cut the RNA.

**Figure 1:**
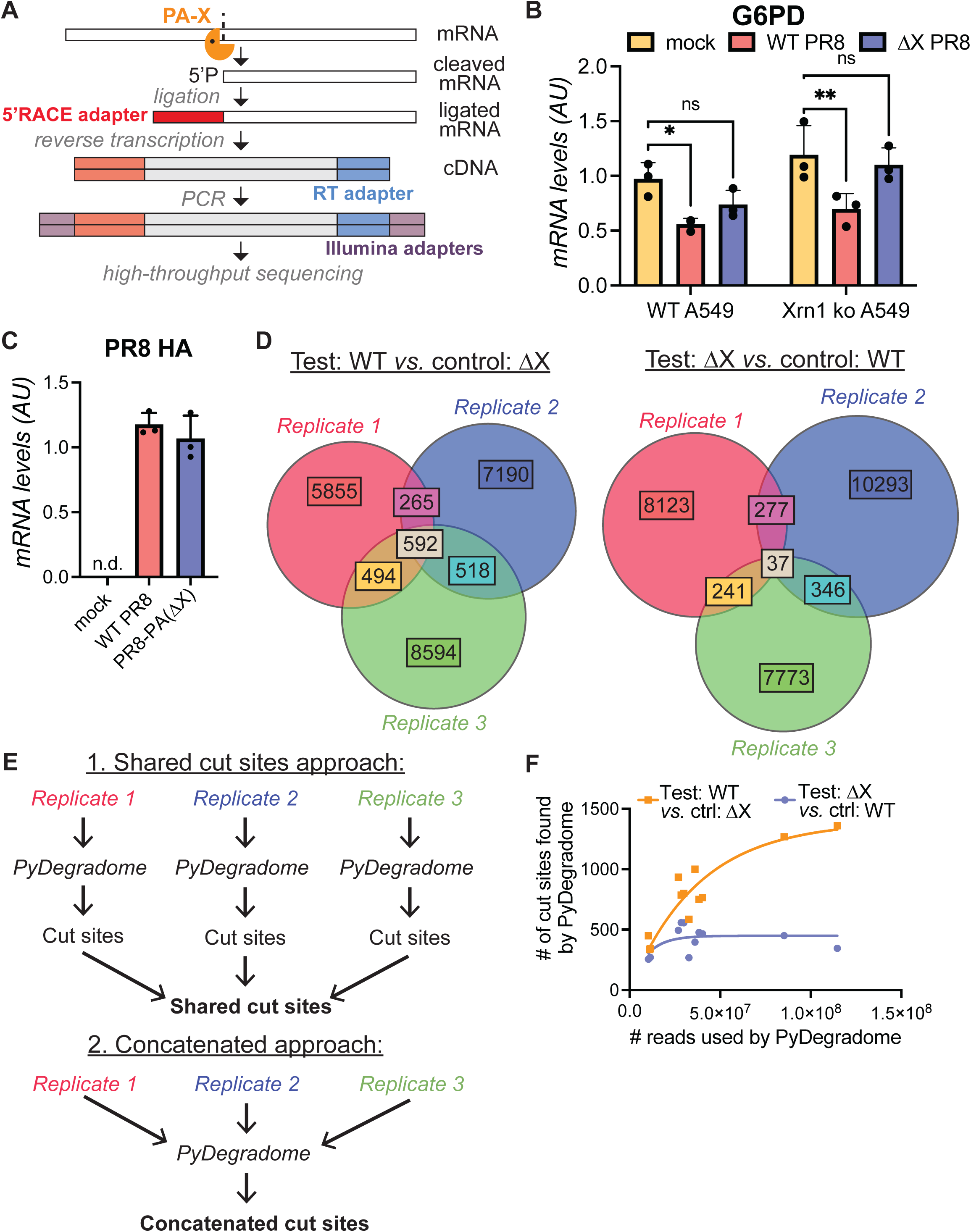
5’ RACE-seq with PyDegradome analysis identifies PA-X cut sites transcriptome-wide. (A) Schematic diagram of 5’ RACE-seq workflow. (B-C) Wild-type (B) or Xrn1 knock out (B-C) A549 cells were either mock infected, infected with WT PR8 or with PR8-PA(ΔX) for 16 hours before RNA extraction. mRNA levels of influenza A virus HA and human G6PD, a PA-X target, were quantified by qRT-PCR, normalized by 18S rRNA levels, and plotted as mean ± standard deviation. AU, arbitrary units; n.d., not defined; ns, not significant; *, p < 0.05; **, p < 0.01; Two-way ANOVA with Dunnett’s multiple comparison test. n = 3 (D) Venn diagrams of the number of cut sites identified by PyDegradome for each replicate. Left diagram shows WT PR8 specific fragments (test: WT PR8, control: PR8-PA(ΔX)), which are likely to be PA-X cut sites; right diagram shows PR8-PA(ΔX) specific fragments (test: PR8-PA(ΔX), control: WT PR8), which are used throughout this study as a negative control. (E) Flow chart of steps used to define PA-X cut sites for further analysis, either through the shared cut sites approach or the concatenated approach. (F) Number of PA-X cut sites (orange squares) or RNA fragments enriched in the PR8-PA(ΔX) infected cells (purple circles) identified by PyDegradome relative to the number of input reads.

Our protocol yielded a high level of background reads, presumably due to the lack of basal RNA degradation in Xrn1 knock-out cells. All libraries had a similar number of reads (**Supplementary Table 1**), and ∼75% of reads mapped to locations that were not shared across samples (**Extended Data Fig. 1D-E**, light blue datapoints). Additionally, the number of reads that started at the same nucleotide across multiple samples was highly correlated (**Extended Data Fig. 1D-E**, black and red datapoints), suggesting common background RNA degradation. However, some locations had higher numbers of reads in infected cells (**Extended Data Fig. 1E**, red datapoints). Some of these fragments could be due to PA-X cleavage. To correct for background degradation and identify locations enriched in WT PR8 compared to PR8-PA(ΔX) infected cells, we used PyDegradome^20^. This pipeline assumes the first read nucleotide after the RNA adapter is the 5’ end of the RNA fragment and counts how many reads within each dataset map to the same 5’ position. These counts are then compared between a test and a control sample, and a Bayesian probability model is used to determine which locations are significantly higher in the test sample. The stringency of cut site definition in PyDegradome can be adjusted based on two user-defined parameters: confidence level (cl) and multiplicative factor (mf) (see methods)^20^. We used loose parameters (cl = 99%, mf = 2) on 16-22 million uniquely mapped reads per sample to find as many cut sites as possible, and identified 592 PA-X cut sites in 517 genes that were shared across three biological replicates (**Fig. 1D**, left, **Supplementary Table 2**). We will refer to this analysis as the “shared” approach (**Fig. 1E**). These sites are likely true PA-X target sites, as a control analysis looking for fragments enriched in the PR8-PA(ΔX) infected cells (i.e. running PyDegradome with PR8-PA(ΔX) infected cells as the test sample and WT PR8 infected cells as the control sample; **Fig. 1D**, right) only returned 37 sites. We considered whether the other non-shared sites we identified could also be PA-X sites, indicating a low specificity for this protein.

However, we found that if we increased the stringency of the analysis (higher cutoff cl and mf values), we increased the percentage of shared sites (**Extended Data Fig. 2A**). This result suggests that at least some of the additional sites are due to either noise or limitations in the pipeline’s ability to distinguish signal from noise. We were also surprised by the relatively low number of RNAs cut by PA-X compared to the >3,000 genes down-regulated by PA-X at steady state levels^4^. We wondered if this low number stemmed from limited sequencing depth and decided to increase it by analyzing the replicates together by concatenating the read files (“concatenated” approach, **Fig. 1E**). While we lose information on the replicability of the cut sites, this method is equally valid from a statistical perspective and has more power. We optimized the parameters by maximizing both the total number of concatenated cut sites detected (**Extended Data Fig. 2B**, red circles) and the percentage of the 592 shared cut sites identified in the concatenated analysis (**Extended Data Fig. 2B**, blue diamonds). Using cl = 99.99% and mf = 2, we identified 1361 cut sites in 1309 genes in WT vs. PR8-PA(ΔX) infected cells (**Supplementary Table 2**), but only 345 cut sites in the control comparison PR8-PA(ΔX) vs. WT PR8.

To further determine if the limited number of identified sites was due to the intensity of the signal and the signal-to-noise ratio, i.e., the number of reads mapping to the cut site in test vs. control samples, we plotted the distribution of the read counts at the cut site for sites identified by one, two or three individual replicates, or by the concatenated approach. The cut sites identified in three replicates or by the concatenated approach had higher reads counts, whereas the ones identified in only one replicate had lower counts (**Extended Data Fig. 2C**), suggesting we may be only identifying the most robust cut sites. Some of the non-shared sites may also be real, but the signal is too low in some replicates to distinguish from background RNA degradation. Considering this result, to determine how much we may benefit from additional sequencing depth, we reanalyzed subsets of our data with the optimal parameters for the concatenated approach, with or without concatenating the read files. We then plotted the number of identified cut sites against the number of input reads. While the number of PA-X cut sites increased with increasing read depth, it plateaued around 100 million reads (**Fig. 1F**, orange squares). In contrast, the number of fragments enriched in the PR8-PA(ΔX) infected cells did not increase with the number of reads (**Fig. 1F**, purple circles). This result suggests that we can use the current analysis without further increasing the sequencing depth, and that we are detecting specific cut sites.

Although PA shares the same endonuclease domain as PA-X and is also active in influenza A virus infected cells, our results suggest we are detecting PA-X-specific cut sites. First, comparing read counts in WT PR8 and PR8-PA(ΔX) infected samples should remove PA cut sites, as PA is equally expressed by both viruses. Indeed, we detected similar levels of PA protein in WT PR8 and PR8-PA(ΔX) infected cells, consistent with previous reports suggesting that PA protein levels are the same or higher in cells infected with PA-X-deficient viruses^3, 5, 6, 13, 24, 25^. Moreover, very few (< 0.2%) of the sites identified by PyDegradome mapped to the first 20 nucleotides of the transcripts, where PA cuts in the context of FluPol (**Extended Data Fig. 2D**). In fact, these 5’ sites were slightly more represented when PA-X was absent, i.e. in the PR8-PA(ΔX)-infected cells vs. mock or WT PR8-infected cells (**Extended Data Fig. 2D**). Thus, the cut sites we identified in WT PR8-infected cells are unlikely to be due to PA.

Given these considerations, we proceeded to further analyze the 1361 cut sites from the concatenated analysis (cl=99.99%/mf=2 parameters on all the reads), which give us a better overview of the PA-X cut site locations transcriptome-wide, as well as the 592 cut sites shared between the three replicates (cl=99%/mf=2 parameters on individual replicates), which have a more biologically stringent cutoff and can inform us on the preferred characteristics of PA-X cut sites (**Supplementary Table 2**).

### PA-X cleavage is driven by RNA sequences

To confirm that PyDegradome identified true PA-X cut sites, we performed classical 5’ RACE (**Fig. 2A**) for 12 sites on RNA from Xrn1 knock-out A549 cells infected with WT or PR8-PA(ΔX), or mock infected, and validated 11, including the 6 examples shown in **Figures 2B** and **Extended Data Fig. 3A**. For this assay, we concluded that the RNA was cut at the predicted location if the RACE PCR amplified a product of the correct size, i.e. the distance between the gene-specific primer and the primer in the ligated adaptor (**Fig. 2A**, blue arrows), only in the WT PR8 infected samples. PCR products were sequenced to confirm the cut site identification. Of note, additional PCR bands were apparent, likely due to basal RNA degradation in Xrn1 knock out cells, but were not specific to the WT PR8 infected cells. We also confirmed by RT-qPCR that WT and PR8- PA(ΔX) samples were similarly infected (**Extended Data Fig. 3B**).

**Figure 2:**
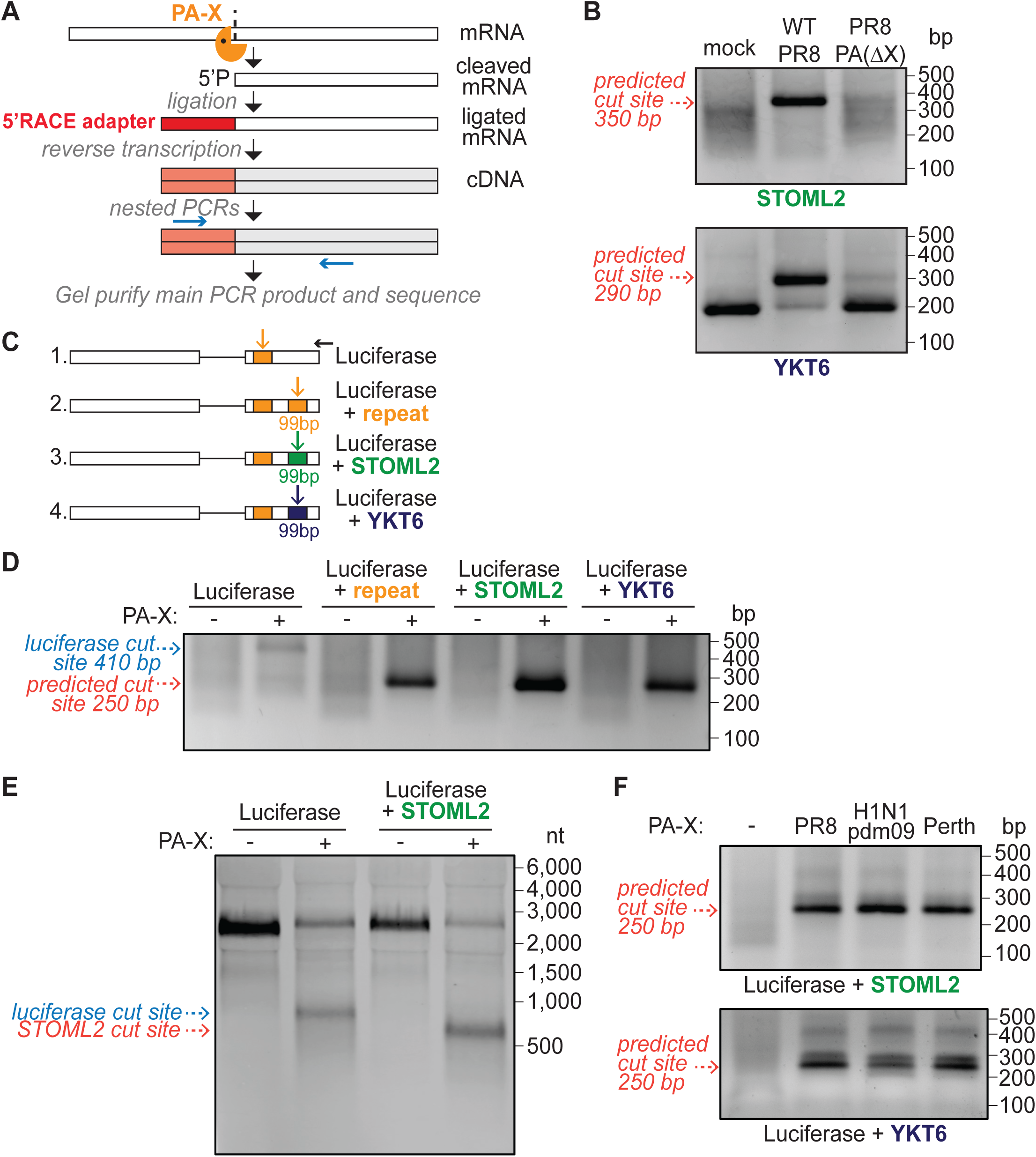
Sequences around the cut sites drive cleavage by PA-X. (A) Schematic diagram of 5’ RACE workflow. (B) Xrn1 knock out A549 cells were infected with WT or PR8-PA(ΔX), or mock infected for 16 hours before RNA extraction. 5’RACE was then performed using primers specific for STOML2 or YKT6, ∼250-300 nucleotides (nt) downstream of the predicted cut sites. The PCR products were run on an agarose gel. The predicted size of DNA bands coming from cut sites identified by PyDegradome are indicated by the red dotted arrows. Background bands can be observed with YKT6 primers at ∼200 bp. These fragments map to TTG//AAC site in exon 7, 93 nt downstream of GCTG cut site, and may represent a constitutive fragment of degradation. (C) Diagram of the luciferase reporters tested. The black horizontal arrow indicates the position of the 5’ RACE PCR reverse primer, vertical arrows indicate the predicted position of the cut sites. (D-F) HEK293T ishXrn1 cells were treated with doxycycline for 3-4 days to induce knock down of Xrn1, then transfected with one of the luciferase reporters in (C), and where indicated, with PA- X from the PR8 (D-F), H1N1pdm09 or Perth influenza strains (F). RNA was extracted to run 5’ RACE (D, F) or northern blotting (E). Expected sizes of DNA/RNA bands coming from cut sites in the introduced target sequences are indicated by the red dotted arrows, while the blue dotted arrow indicates the size of the original luciferase cut site fragment. For all DNA gels, the DNA bands were purified and sequenced to confirm their identities. Gel and northern blot images are representative of 3 experiments.

To test whether RNA sequences drive PA-X cleavage, we introduced 99 base pairs (bp) surrounding several PA-X cut sites into a luciferase reporter that contains an intron and is down- regulated by PA-X^4, 26^ (**Fig. 2C**, constructs 3 and 4). We transfected this reporter with or without PR8 PA-X into human embryonic kidney (HEK) 293T cells expressing inducible shRNA against Xrn1 (ishXrn1), which were pre-treated with doxycycline to knock down Xrn1 (**Extended Data Fig. 3C**). As negative and positive controls, we used the original luciferase reporter with no inserted sequence, which has an “endogenous” cut site (**Fig. 2C**, construct 1, orange rectangle) and a construct in which we duplicated the 99 bp sequence around this original cut site (**Fig. 2C**, construct 2). Introducing extra sequences into the luciferase reporter did not interfere with its expression and down-regulation by PA-X (**Extended Data Fig. 3D**), but added a new PA-X cut site (red dotted arrows, **Fig. 2D**, **Extended Data Fig. 3E**). For the negative control, we detected fragments from the original cut site instead (blue dotted arrows, **Fig. 2D**, **Extended Data Fig. 3E**). We also confirmed these results using northern blotting, which detected single fragments coming from the luciferase and luciferase + STOML2 reporters (**Fig. 2E**). While RACE could selectively amplify some fragments, northern blotting should allow us to see multiple RNA fragments if present. Therefore, the results in **Fig. 2E** support our model that PA-X cleaves RNAs at discrete sites. Overall, these results suggest that the sequences around the cut sites are sufficient to drive cleavage by PA-X.

Importantly, the sequences we identified only drive cleavage by PA-X. We detected no cleavage upon expression of the PA-X D108A catalytic mutant, a PA(fs) mutant that makes PA but not PA-X due to a decrease in frameshifting^4^, or other viral host shutoff endoRNases (**Extended Data Fig. 4A-B**). Conversely, these sequences also elicited cleavage by PA-X from the influenza A/Perth/16/2009 (H3N2) (henceforth “Perth”) and A/Tennessee/1-560/2009 (H1N1) (henceforth “H1N1pdm09” for 2009 pandemic H1N1) viruses, representative of currently circulating H3N2 and H1N1 A strains (**Fig. 2F**, **Extended Data Fig. 4C**). We also saw conserved cleavage specificity during infection with Perth and H1N1pdm09 viruses (**Extended Data Fig. 4D**). The PA-X proteins from PR8, Perth and H1N1pdm09 (**Extended Data Fig. 4E**) are a good representation of PA-X diversity across human influenza A strains, as they represent three different subtypes of influenza A viruses and have different C terminus lengths (41 aa for H1N1pdm09 *vs.* 61 aa for PR8 and Perth) and half-lives^27^. These experiments confirm that 5’ RACE-seq and PyDegradome identified bona fide PA-X cut sites transcriptome-wide and indicate that PA-X cleavage across multiple strains is sequence specific.

### PA-X preferentially cleaves RNAs at a specific sequence and structure

Because the sequences surrounding PA-X cut sites were sufficient to trigger cleavage, we wondered whether conserved RNA features guided cleavage. We saw a depletion in adenosines and an enrichment in GCUG at the cut site, and a modestly enriched guanidine stretch downstream of the cut site (**Fig. 3A** 592 sites from shared analysis, **Extended Data Fig. 5A** 1361 cut sites from concatenated analysis). These features were specific to PA-X, as we found different patterns in fragments specific to PR8-PA(ΔX) infected cells, our control comparison (**Fig. 3B**, **Extended Data Fig. 5B**). Interestingly, most PA-X cut sites contained GCUG or a similar tetramer, although the location of the cut within the tetramer varied (**Fig. 3C**, **Extended Data Fig. 5C**). The first two nucleotides in GCUG were the most important for PA-X cleavage (**Fig. 3D**, **Extended Data Fig. 5D**).

**Figure 3:**
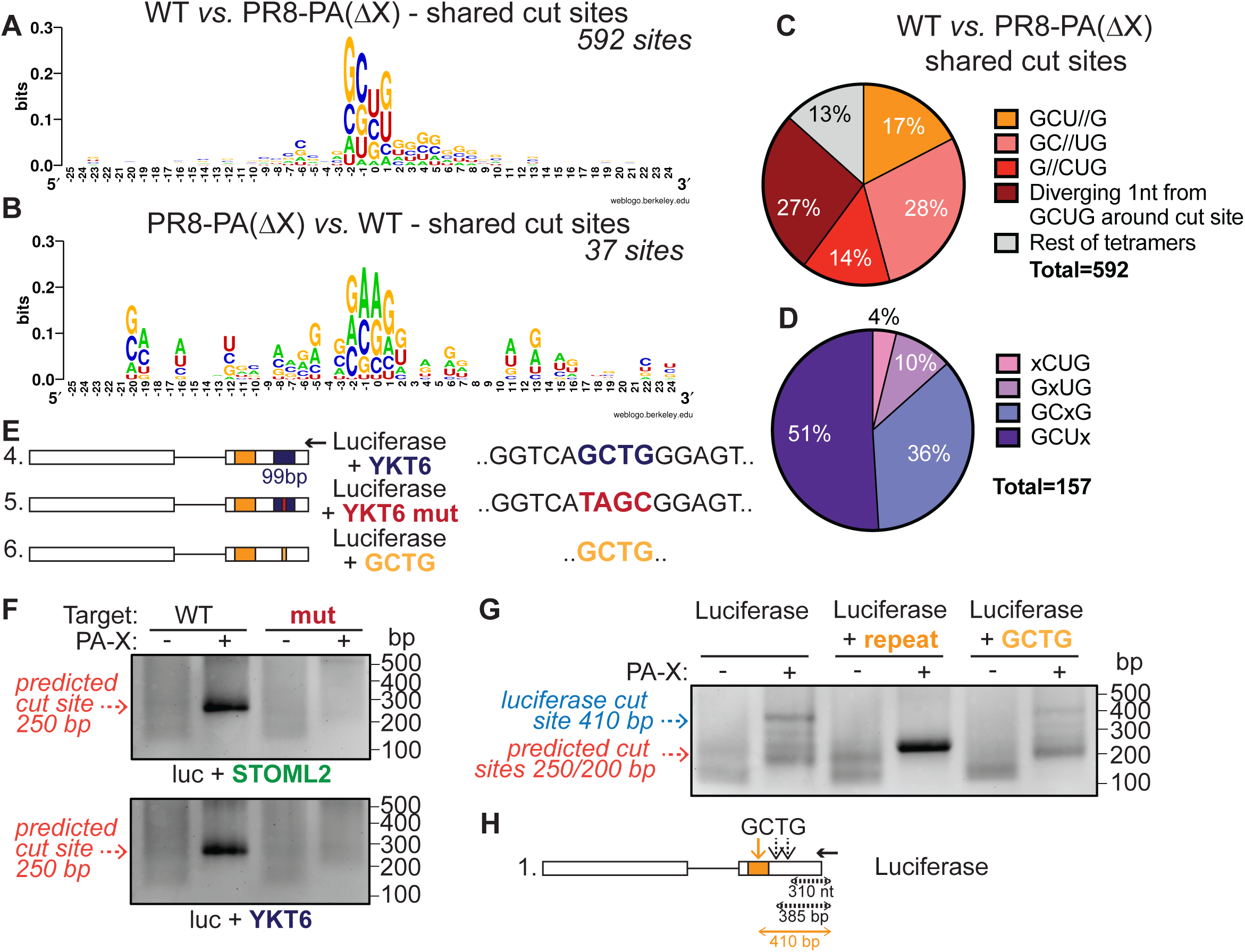
PA-X preferentially cleaves RNAs at GCUG tetramers. (A-B) WebLogo^70^ representation of base enrichment around PA-X cut sites (test: WT PR8 vs. control: PR8-PA(ΔX), A) or around control sites enriched in the PR8-PA(ΔX) sample (test: PR8-PA(ΔX) vs. control: WT PR8, B) for sites predicted by PyDegradome using the shared cut sites approach. (C) Percentage of PA-X cut sites containing GCUG or a tetramer with one nucleotide difference from GCUG, for sites identified by PyDegradome using the shared cut sites approach. // indicates the location of the cut, i.e. GCU//G indicates that PA-X cuts between the U and the G. (D) Further breakdown of the PA-X cut sites containing a tetramer with one nucleotide difference from GCUG around the cut site (marked by an x). (E) Diagram of the luciferase reporters tested. The black arrow indicates the position of the 5’ RACE PCR reverse primer used. (F-G) HEK293T ishXrn1cells were treated with doxycycline for 3-4 days to induce knock down of Xrn1, then transfected with luciferase reporters containing 99 bp insertions from the indicated genes with or without the GCTG à TAGC mutation shown in E (F), a repeat of the luciferase cut site (G), or the GCTG tetramer alone (G), with or without PR8 PA-X. RNA was extracted to run 5’ RACE. Expected sizes of DNA bands coming from cut sites in the introduced target sequences are indicated by the red dotted arrows (∼250 bp for luciferase + STOML2, luciferase + YKT6 and luciferase + repeat, ∼200 bp for luciferase + GCUG), while the dotted blue arrow indicates the size of the original luciferase cut site fragment. DNA bands were purified and sequenced to confirm their identities. Gel images are representative of 3 experiments. (H) Diagram of the positions of several GCTG tetramers in the luciferase reporter. The orange vertical arrow indicates the original luciferase cut site and the black vertical dotted arrows additional GCTG sites that are not clearly cleaved by PA-X. Black horizontal arrow indicates location of 5’ RACE reverse primer, and lengths underneath diagram indicate the expected length of 5’ RACE PCR products each cut site would generate.

To test the importance of the GCUG tetramer, we mutated the cut site GCTG to TAGC in the luciferase reporters with the 99 bp insertions from BCAP31, STOML2, TUBA1B and YKT6. (**Fig. 3E**, construct 5). This mutation did not impact expression of the reporters, or their down-regulation by PA-X since PA-X still cleaves the original luciferase sequence **(Extended Data** **Fig. 5E**). However, the mutation prevented efficient cleavage by PA-X within the inserted sequence (**Fig. 3F**, **Extended Data Fig. 5F**). However, introduction of GCTG alone (**Fig. 3E**, construct 6) did not result in efficient cleavage (**Fig. 3G**). Thus, GCUG is necessary but not sufficient for PA-X cleavage, consistent with PA-X not cleaving at every GCUG. For example, there are GCUG tetramers 17 and 92 nucleotides downstream of the original cut site in the luciferase mRNA (**Fig. 3H**, black vertical dotted arrows), but we do not see evidence of robust cleavage by PA-X at these locations.

While GCUG alone was not sufficient, the minimal sequence required for PA-X cleavage varied depending on the gene tested (**Fig. 4A**). Whereas inserting 15 bp was sufficient for PA-X cleavage of the STOML2, BCAP31 and SLC7A5 cut sites, more than 27 bp were required for the YKT6 sequence (**Fig. 4B**, **Extended Data Fig. 6A**). Given the lack of additional enriched motifs (**Fig. 3A**, **Extended Data Fig. 5A**), we tested the potential role of RNA secondary structures. We predicted the RNA structure around the shared PA-X cut sites with LinearFold^28^, using either the CONTRAfold v2.0 machine-learning model^29^ (LinearFold C) or the Vienna RNAfold thermodynamic model^30, 31^ (LinearFold V). In both models, PA-X cut sites (but not control sites enriched in PR8-PA(ΔX) infected cells) were preferentially found in the loops of RNA hairpin structures (**Fig. 4C**). To test whether the hairpin was required for cleavage, we disrupted the predicted hairpin structures in the YKT6 51 bp and the STOML2 15 bp insertion constructs (**Fig. 4A**). We mutated select C or G bases to break strong G-C bonds in the stems (**Fig. 4D-E**, *WT*, red arrows), leading to very different predicted structures, with the GCUG cut site in bulge or paired regions (**Fig. 4D-E**, *GC unpaired mutant*). As a control, we additionally mutated the complementary G or C bases to recreate the G-C bonds and repair the structures (**Fig. 4D-E**, *GC repaired mutant*, blue arrows). Consistent with PA-X cut site structure predictions, disrupting the hairpin prevented efficient PA-X cleavage within the YKT6 and STOML2 sequences. The reporter was instead cleaved at the original luciferase cut site location (**Fig. 4F-G**, blue dotted arrow). Conversely, repairing the hairpin structure restored efficient cleavage by PA-X within the inserted fragment despite the mutated surrounding sequence (**Fig. 4F-G**, red dotted arrow). Selective PA- X cleavage of GCUG sequences in a hairpin loop was also seen using the same strategy during infection (**Extended Data Fig. 6B**). Thus, PA-X preferentially cleaves RNAs at GCUG tetramers located within hairpin loops, which reveals a new layer of RNA targeting specificity by PA-X.

**Figure 4:**
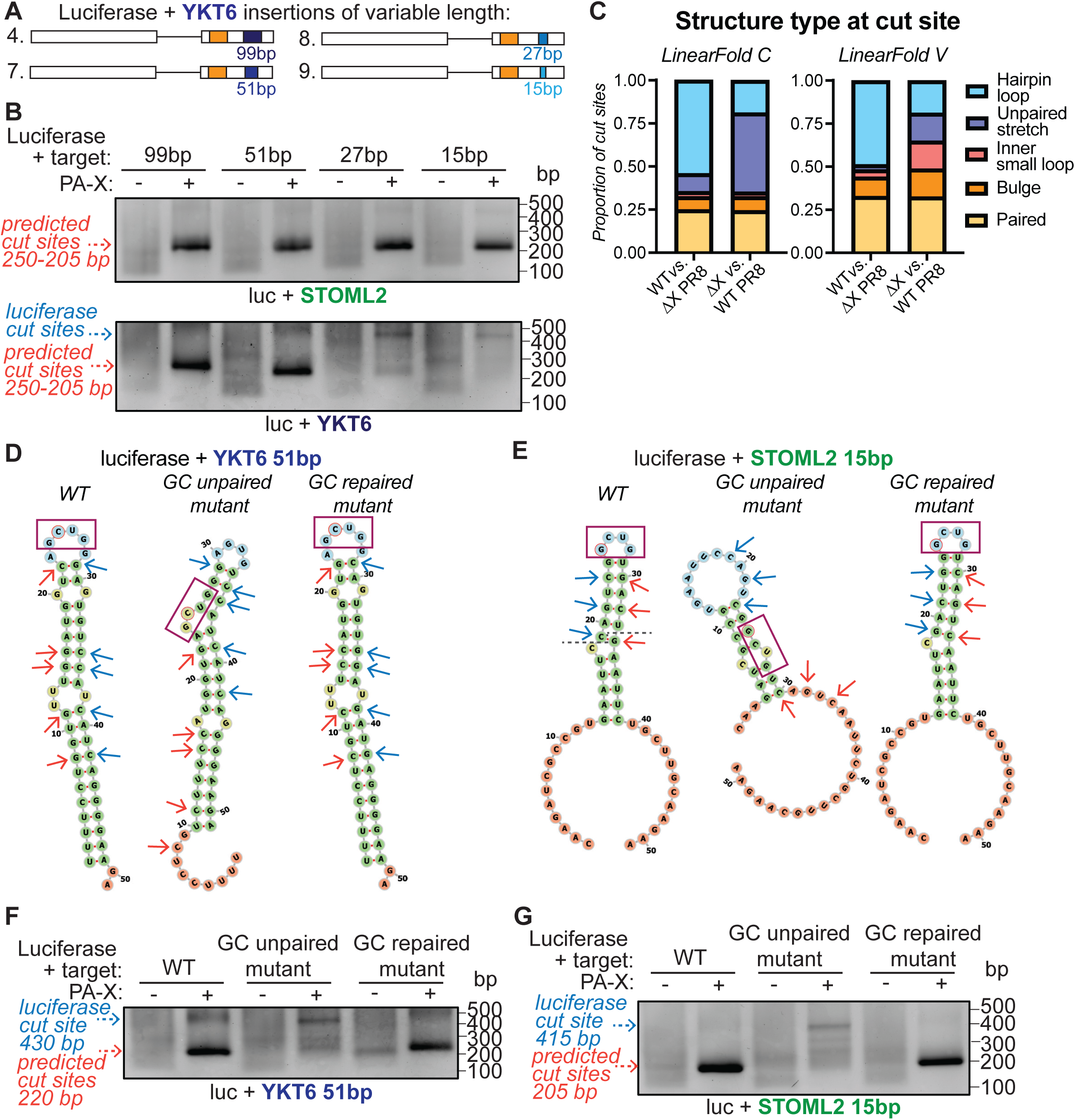
PA-X preferentially cleaves RNA within hairpin loop structures. (A) Diagram of the luciferase reporters tested. (B, F, G) HEK293T ishXrn1cells were treated with doxycycline for 3- 4 days to induce knock down of Xrn1, then transfected with luciferase reporters containing insertions of the indicated lengths from the YKT6 or STOML2 genes (B), with or without the mutations indicated in D-E (F-G), with or without PR8 PA-X. RNA was extracted to run 5’ RACE. Expected sizes of DNA bands coming from cut sites in the introduced target sequences are indicated by the red dotted arrows (∼250 bp for 99 bp constructs, ∼220 bp for 51 bp constructs, ∼210 bp for 27 bp constructs and ∼205 bp for the 15 bp constructs). Blue dotted arrows indicate the size of the original luciferase cut site fragments (∼430 bp for 51 bp constructs, ∼420 bp for 27 bp constructs and ∼415 bp for 15 bp constructs). DNA bands were purified and sequenced to confirm their identities. Gel images representative of 3 experiments. (C) Predicted RNA secondary structures of the 99 bp sequence around PA-X cut sites (WT PR8 vs. PR8 PA(ΔX), n = 592) or around control sites enriched in the PR8-PA(ΔX) sample (PR8 PA(ΔX) vs. WT PR8, n =37) (shared cut sites approach). Structures were predicted using the CONTRAfold v2.0 machine- learning model (LinearFold C) or the Vienna RNAfold thermodynamic model (LinearFold V). (D-E) Diagram of the LinearFold C predicted structures for the 51 bp YKT6 sequence (D) or the 15 bp STOML2 and surrounding luciferase sequences (E). Left = structures for the WT sequences, middle = structures for sequences with mutations in the nucleotides indicated by the red arrows (GC unpaired mutant), right = structures for sequences with mutations in the nucleotides indicated by both the red and blue arrows (GC repaired mutant). The GCUG cut sites are indicated by the purple boxes. PA-X cuts these sequences after the “C” or “G” circled in red. In E, the dark grey dotted line on the WT diagram indicates where the STOML2 sequence ends and the luciferase surrounding sequence begins.

### PA-X cleaves RNAs within exons

Since PA-X activity is linked to splicing^4^, we wondered whether PA-X targets pre-mRNAs or mature mRNAs. Interestingly, PA-X may preferentially cleave RNA within exons, as cut sites were almost exclusively found within exons (**Fig. 5A**) although 12-16% of reads mapped to introns (**Fig. 5B**, p < 0.001, Chi-Square Test for Goodness of Fit with degree of freedom 1). In contrast, 18% of PR8-PA(ΔX) specific fragments from the concatenated analysis mapped to introns (**Extended Data Fig. 7A**, bottom). While only 3% of PR8-PA(ΔX) fragments from the shared analysis mapped to introns, this may stem from the very low number of sites (37) found with this method (**Extended Data Fig. 7A**, top).

**Figure 5:**
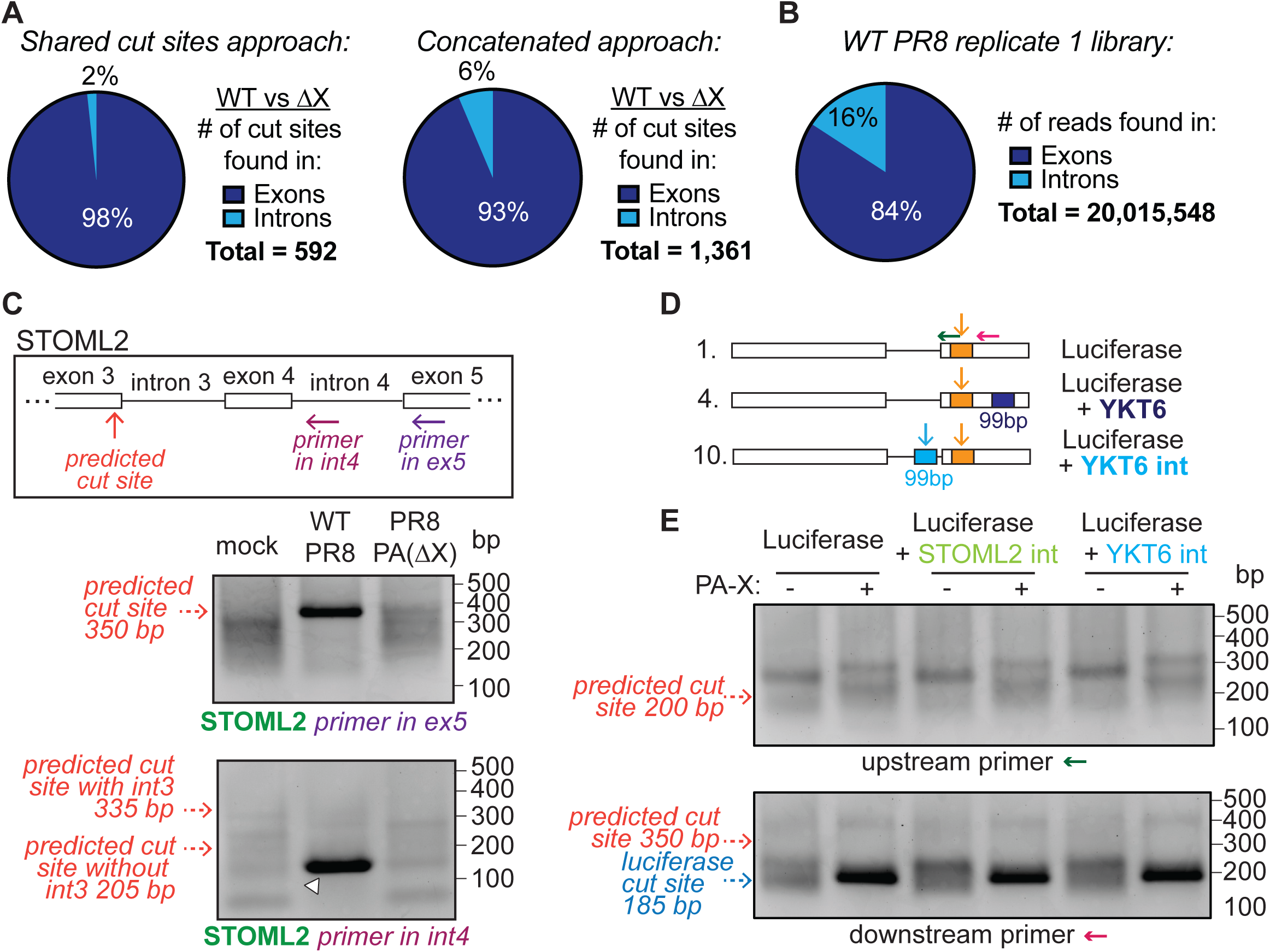
PA-X preferentially cleaves RNAs within exons. (A-B) Percentage of PA-X cut sites found within introns or exons, for sites identified by PyDegradome using the shared cut sites approach (A, left) or the concatenated approach (A, right), compared to the percentage of reads found in exons vs. introns (B). The reads used in B are from WT PR8 replicate 1 as an example, but other samples have similar read distribution. (C) Xrn1 knock out A549 cells were infected with WT PR8 or PR8-PA(ΔX), or mock infected. 5’RACE was performed and PCR products were run on an agarose gel. The box shows a diagram of the STOML2 gene and the positions of the reverse primers for 5’ RACE. The top gel is the same gel as Fig. 2B and is included for comparison. A reverse primer in exon 5 was used for the PCR, and the red dotted arrows indicate size of fragment originating from previously validated cut site. The bottom gel shows products obtained using a reverse primer in intron 4, and red dotted arrows indicate the predicted size of PCR products that would appear if PA-X cleaved unspliced pre-mRNAs. White arrowhead indicates a fragment mapped to exon/intron 4 junction. Gel images are representative of 3 experiments. (D) Diagram of the luciferase reporters tested. Green and magenta arrows indicate positions of 5’ RACE PCR reverse primers, vertical arrows indicate location of predicted cut sites. (E) HEK293T ishXrn1 cells were treated with doxycycline for 3-4 days to induce knock down of Xrn1, then transfected with luciferase reporters containing 99 bp insertions from the STOML2 or YKT6 gene in the intron, as shown in (D), with or without PR8 PA-X. RNA was extracted and used to run 5’ RACE. PCR products were separated on an agarose gel. The top gel represents products obtained using green arrow primer from D and the bottom gel using magenta arrow primer from D. Blue dotted arrow indicates size of PCR products originating from the original luciferase cut site, red dotted arrows indicate the predicted sizes of PCR products that would originate from the sequence inserted in the introns. Gel images representative of 3 experiments.

Our 5’ RACE validation experiments (**Fig. 2B**, **Extended Data Fig. 3A**) also suggested that PA-X cleaves after splicing, as the RACE primers spanned exon-exon junctions but only amplified spliced fragments (**Fig. 5C**, **Extended Data Fig. 7B**, dark purple arrows). However, this could be an artifact of PCR, which may favor the amplification of smaller spliced fragments over longer non-spliced ones. We thus repeated the 5’ RACE PCRs using reverse primers that should amplify cleaved unspliced pre-mRNAs (**Fig. 5C**, **Extended Data Fig. 7B**, light purple primers). However, we did not detect fragments of expected sizes specifically in WT PR8 infected cells, but instead saw background fragments, including some that mapped to the exon/intron junction (**Fig. 5C**, **Extended Data Fig. 7B**, white arrowheads). This result suggests that PA-X cleaves RNAs after splicing of the neighboring introns.

As an additional test, we cloned the previously validated STOML2 and YKT6 99 bp cut site sequences inside the luciferase reporter intron (**Fig. 5D**, construct 10), which did not affect mRNA expression or splicing (**Extended Data Fig. 7C**). However, 5’ RACE did not amplify bands that would correspond to cuts within the intron (**Fig. 5E**, top gel, see dark green arrow in **Fig. 5D** for position of the primer). The closest band at ∼250 bp was also present in the luciferase construct without the STOML2 or YKT6 cut site sequences. Like in other cases where no additional cut site was introduced (**Fig. 3G**, **4B**, **4F-G**), we could still detect the original luciferase cut site using a reverse primer positioned further downstream (**Fig. 5E**, bottom gel, see magenta arrow in **Fig. 5D** for the position of the primer). We also mutated the original luciferase cut site GCUG sequence to UAGC to reduce cleavage at this site (**Extended Data Fig. 7D**, construct 11). However, we still saw no PCR product matching a cut site in the luciferase intron (**Extended Data Fig. 7E**). These results suggest that the STOML2 and YKT6 99 bp sequences are sufficient for PA-X cleavage only when located within an exon and that PA-X cuts mRNAs within exonic sequences after splicing of at least the neighboring intron.

### GCUG tetramers are more abundant in the human than influenza transcriptome

Finally, we wondered why PA-X has evolved these preferred cleavage characteristics, and if this could lead to targeting of specific RNAs. Interestingly, GCUG is one of the most abundant tetramers in the human transcriptome (**Extended Data Fig. 8A**) and is found at twice the frequency in the transcriptome compared to the human genome, which includes the intronic sequences (**Fig. 6A**). Given that mRNAs cut by PA-X do tend to have more GCUG tetramers (**Fig. 6B**), PA-X may have evolved these cleavage characteristics to preferentially target the host transcriptome.

**Figure 6:**
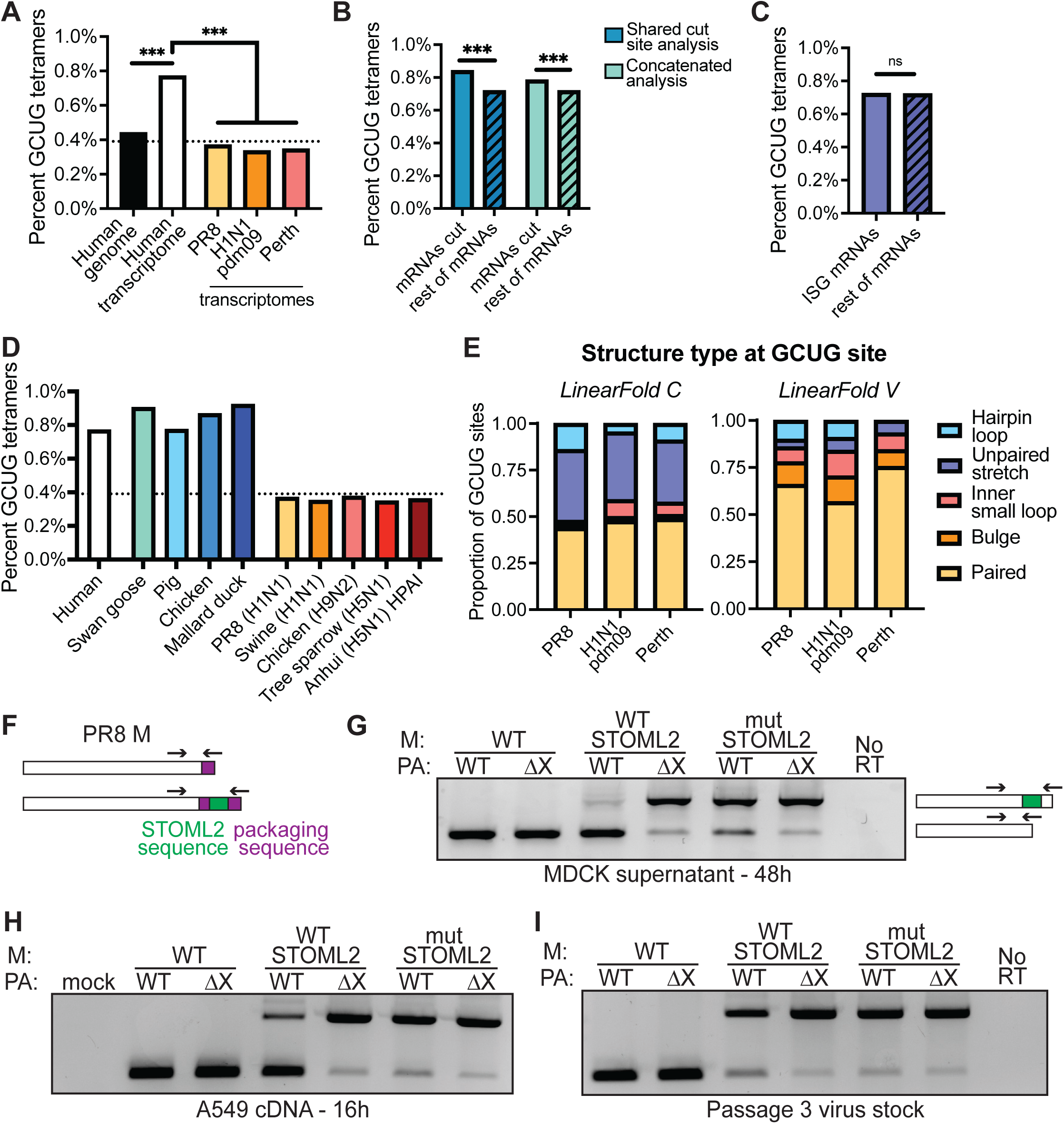
PA-X likely cleaves GCUG sequences to preferentially target host over viral mRNAs. (A-D) The percentage of GCUG tetramers in each indicated sequence was calculated by counting the number of GCUG tetramers and dividing it by the total number of tetramers in the sequence (i.e. length of the sequence minus 3). ISG: interferon stimulated genes; ns = not significant, *** = p < 0.001, Chi-Square Test for Goodness of Fit with degree of freedom 1. The black dotted lines in (A) and (D) represent the average tetramer abundance expected by chance (i.e. 1/256). Transcriptomes in (D) are from Anser cygnoides (swan goose), Sus scrofa (pig), Gallus gallus (chicken), Anas platyrhynchos (duck), A/swine/Guangdong/2722/2011 (H1N1), A/chicken/Pakistan/UDL/01/2008 (H9N2), A/tree sparrow/Jiangsu/1/2008 (H5N1) and A/Anhui/1/2005 (H5N1) highly pathogenic avian influenza (HPAI) (E) Predicted RNA secondary structures of the 99 nt sequence around GCUG tetramers in the indicated influenza transcriptomes. Structures were predicted using the CONTRAfold v2.0 machine-learning model (LinearFold C) or the Vienna RNAfold thermodynamic model (LinearFold V). (F) Diagram showing the location of the insertion of the STOML2 51 bp cut site sequence (green rectangle) in the PR8 M segment after the end of the M2 coding sequence. The full M segment packaging signal (purple rectangle) was positioned after the STOML2 sequence to make sure the modified segment is packaged inside the virion, resulting in a duplication of some sequences including the end of the M2 coding region. The position of primers to amplify sequences at the end of the M segment is marked by arrows. The reverse primer only binds the full packaging sequence at the very end of the engineered M-STOML2 fusion segment. (G-I) Viral RNA was extracted from 48-hour supernatants from infected MDCK cells from the experiment shown in Extended Data Fig. 9C (G), Xrn1 knock out A549 cells 16 hours post-infection (H), or viral stocks (I), for viruses harboring no insert, inserted WT STOML2 or mutant STOML2 sequences, either in the WT PR8 or PR8- PA(ΔX) background. Mutant STOML2 sequences contained the GCUG à UAGC mutation that prevents PA-X cleavage (Fig. 3). The RNA was then reverse-transcribed to cDNA and PCR amplified using primers located on either side of the STOML2 sequence (black arrows in F-G) to determine whether the STOML2 sequence was retained. Gel images are representative of 3 experiments with separate virus rescues.

It is unclear if GCUG specificity has any role in targeting specific groups of host mRNAs. For example, the mRNAs for the antiviral interferon stimulated genes (ISGs) do not contain more GCUG tetramers than other mRNAs (**Fig. 6C**). However, GCUG specificity may contribute to virus vs. host discrimination. Indeed, whereas GCUG is abundant and overrepresented in the human transcriptome, it is far less abundant in the mRNAs of three influenza A virus strains, PR8 H1N1, H1N1pdm09 and Perth H3N2 (**Fig. 6A, Extended Data Fig. 8B**) and is underrepresented relative to other tetramers (**Extended Data Fig. 8C-E**). GCUG levels are also higher in the cellular transcriptomes of common influenza hosts such as pigs, chickens, geese and ducks, than in the transcriptomes of representative influenza A virus strains isolated from these animals (**Fig. 6D**). The differential GCUG representation is consistent with our finding that PA-X activity does not down-regulate influenza transcripts^13^, and points to GCUG specificity as a way to distinguish host vs. viral mRNA. In support of this idea, the percentage of GCUG is higher in the negative RNA strand (i.e. the viral genomic RNA) than in the positive RNA strand (i.e. the viral mRNA) for almost all segments (**Extended Data Fig. 8B**). Additionally, LinearFold structure predictions suggested that GCUG tetramers in influenza transcripts are enriched in paired structures that may not be accessible to PA-X (**Fig. 6E**). We also tested whether PA-X cuts GCUG tetramers in the viral transcriptome predicted to be within hairpin loops. We detected faint bands that mapped close to the GCUG tetramers in the case of PR8 (**Extended Data Fig. 9A**), but no cut sites for H1N1pdm09 (**Extended Data Fig. 9B**), suggesting that viral mRNAs are cut very inefficiently, if at all. These findings suggest that PA-X may have evolved to target GCUG sequences in hairpin loops to avoid degrading influenza mRNAs while preferentially targeting the host transcriptome.

To test this idea experimentally, we inserted the STOML2 51 bp cut site sequence inside the PR8 M segment (segment 7) (**Fig. 6F**). We picked a segment that is spliced because we previously found that PA-X activity is linked to splicing^4^. As a control, we inserted the STOML2 51 bp sequence with GCUG mutated to UAGC, as this mutation prevented efficient PA-X cleavage (**Fig. 3E-F, Extended Data** **Fig. 5F**). Initial viral growth assays did not reveal any differences in viral growth between WT PR8 and PR8-PA(ΔX) with or without the two inserts (**Extended Data Fig. 9C**). However, when we purified the viral RNA present in the supernatant 48 hours post infection, we noticed that the inserted STOML2 sequence had been lost when PA-X was present (i.e. in the WT PR8 background) (**Fig. 6G**). In contrast, it was retained in the absence of PA-X (i.e. in the PR8-PA(ΔX) background) (**Fig. 6G**). Moreover, the mutated STOML2 sequence that is not cleaved by PA-X was still present in both WT PR8 and PR8-PA(ΔX) backgrounds (**Fig. 6G**). Infecting A549 cells overnight also lead to loss of the STOML2 sequence insertion (**Fig. 6H**), even though the STOML2 sequence was present in the viral inoculum (**Fig. 6I**). We rescued the STOML2 recombinant viruses three separate times and in every replicate the inserted STOML2 sequence was selectively lost in the presence of PA-X. These results suggest that during viral propagation, viruses that have lost the PA-X cleavage site have an advantage for growth, i.e that a preferred PA-X cut site inside a spliced viral segment is detrimental for the virus. This is only true when PA-X is present, confirming that the insertion itself is not deleterious. Overall, these results are consistent with PA-X evolving specific cleavage characteristics to preferentially target host over viral mRNAs.

## Discussion

Studies of viral host shutoff endoRNases have mostly relied on RNA steady-state levels as a proxy for RNA degradation. However, widespread RNA degradation can have downstream effects on other aspects of RNA metabolism^1^. Steady-state level analysis may thus preclude a clear understanding of how these enzymes target RNAs and which RNAs they degrade. Therefore, we examined RNA degradation directly by identifying PA-X cut sites transcriptome-wide and analyzing their characteristics in infected cells. We found that PA-X preferentially cuts RNAs at GCUG or similar tetramers located within hairpin loops, and likely after splicing. This sequence/structure preference is conserved across PA-X from multiple influenza strains. The preferred cleavage sequence is more frequent in human transcripts than in intergenic regions or introns, and most importantly, than in influenza mRNAs. Moreover, PA-X prevents the growth of a recombinant virus carrying a preferred PA-X target sequence. Collectively these results suggest that influenza A virus has evolved a self vs. non-self discrimination mechanism, similar to cellular immune mechanisms that distinguish between host and viral nucleic acids, to induce host shutoff across a broad range of host RNA without hampering its own protein expression.

A previous *in vitro* study of PA-X cleavage activity did not identify sequence specificity^12^, likely because these characteristics can be missed *in vitro* without prior knowledge, highlighting the advantage of our technique. The study did find that PA-X preferentially cuts single-stranded RNA, likely explaining the preference for single-stranded loops. Interestingly, while PA and PA- X share the same RNase domain, they seem to have different cut site specificity^12^, probably due to their different C-terminal domains. Studies that investigated which host RNAs are cap-snatched by PA have found little sequence specificity^32–36^, suggesting that the location of PA cleavage is likely governed by the distance between the cap binding site in the polymerase basic 2 (PB2) subunit of FluPol and the PA active site. We did not see cap-snatching fragments in our dataset, likely because mRNA decapping in the nucleus leads to RNA degradation by the Xrn2 RNase, which is still present in our cells^37^. It would thus be interesting to repeat the experiments in Xrn2 knock out/down cells, and also test whether Xrn2 has an effect on PA-X fragments in the nucleus. A testimony to the importance of the sequence and structure preference of PA-X is that it is conserved across multiple influenza A strains (**Fig. 2F, Extended Data** **Fig. 4C-D**). This is particularly interesting as PA-Xs from different strains have differences in host shutoff activity^38–41^. However, the 5’ RACE PCRs make this method very sensitive but not very quantitative. Therefore, it is difficult to say whether there are quantitative differences in efficiency of cleavage between strains. Nonetheless, the specific sequence and structure that is preferred by PA-X (**Fig. 3-4**) is abundantly found in host mRNAs. Recognizing a ubiquitous motif may be a strategy to target a broad range of host mRNAs. Supporting this idea, we reported a similar specificity for a degenerate recognition motif for another viral RNase, SOX from KSHV^20^. Additionally, both PA- X and SOX, as well as the human endoRNase MCPIP1/Regnase-1^42^, prefer cutting RNA within hairpin loop structures. Targeting exposed loops may be an easy way for endoRNases to access and cleave single-stranded RNA. Studying the cut site characteristics of other viral and human endoRNases will be important to test these ideas.

Two key questions in host shutoff are how the molecular activity of host shutoff factors is linked to immunomodulation and how it is regulated to prevent inhibition of viral gene expression. While we did not uncover a relationship between PA-X cleavage specificity and immunomodulation, we found that it may contribute to PA-X ability to spare viral mRNAs from degradation. Our analysis indicates that influenza A virus mRNAs contain few GCUG tetramers, mostly in paired regions (**Fig. 6**). This result is exciting as it provides an additional mechanism for virus vs. host discrimination by PA-X. Interestingly, most viral genomic RNAs have a higher GCUG percentage than their complementary mRNAs. This result may indicate that there is less pressure to decrease the GCUG percentage in the influenza genome, perhaps because the genomic segments are associated with NP, which could protect them from cleavage^43^. Conversely, GCUG is one of the most abundant tetramers in the human transcriptome (**Extended Data Fig. 8**). PA-X could have evolved this optimal target sequence to destroy as many host transcripts as possible without also targeting anything unnecessary. Consistent with this idea, inserting a sequence targeted by PA-X in one of the viral segments is detrimental to the virus only in the presence of PA-X (**Fig. 6F-I**). Therefore, we propose that this targeting specificity may act as a self vs. non-self discrimination mechanism for influenza A virus. Self/non-self discrimination is common in immune responses, as the cell needs to discriminate its own nucleic acids from those of invading pathogens to prevent inappropriate activation of immune and inflammatory signals. Common discrimination features include RNA modification, double-stranded RNA structures and DNA methylation. However, sequences can also be important. For example, the ZAP protein targets viral RNAs for degradation based on the presence of CG dinucleotides^44^ and the Toll-like receptor TLR9 has preferred target sequences^45^. Our results suggest that influenza A virus is using a similar process in reverse, identifying cellular RNAs as non-self due to the abundance of specific sequences.

The fact that we only found 500-1,400 cut sites through 5’ RACE-seq but that most host RNAs are down-regulated upon PA-X expression^4^ is intriguing. Technical aspects of the method could limit our ability to capture more sites, even though our follow-up analyses suggest the cut sites we did find were true PA-X target sites and could reveal preferred sequence characteristics. Nonetheless, we cannot conclude that these are the only RNAs directly cut by PA-X. Indeed, we detected many sites that appeared to not be shared between replicates. These sites tended to have lower read counts than the PA-X cut sites that were shared by all replicates or that were detected by the concatenated approach (**Extended Data Fig. 2C**). There are several possibilities as to what may cause this result. The sites may not be PA-X cut sites at all but may be selected by PyDegradome due to limitations in distinguishing signal from noise. Alternatively, PA-X may cut different RNAs with varying efficiency depending on how close sequences match the preferred cut characteristics. Thus, these additional sites could be cut by PA-X at lower efficiency. Lastly, we did find that the transcripts that harbor PA-X cut sites are expressed at higher levels (data not shown), suggesting these additional sites may also be real PA-X cleavage locations that raise to the level of significance only in some replicates due to low gene expression. In the future, we will continue to refine our pipeline to improve on these issues. It remains possible, however, that despite these limitations we are indeed capturing the majority of PA-X cut sites in a cell. This possibility would imply that PA-X may not need to cleave thousands of RNAs to trigger widespread down-regulation of RNAs. Rapid cellular RNA degradation due to expression of other viral RNases, infection with the murine gammaherpesvirus 68 (MHV68) and apoptosis causes a secondary inhibition of transcription^21, 46, 47^. It is thus unclear how much of the widespread depletion in host RNAs during influenza infection and PA-X expression is due to RNA degradation *vs.* decrease in transcription. A decrease in transcription has been observed during influenza infection, although it may be partly driven by another influenza protein, non-structural protein 1^48, 49^. Future studies will need to decouple RNA degradation from transcription. It would also be interesting to determine if specific RNAs need to be degraded by PA-X to induce host shutoff, or if the number of degraded RNAs and/or the speed of degradation determines transcription inhibition. Finally, inhibition of transcription could also dampen the induction of antiviral RNAs when PA-X is present^4^, even though few antiviral RNAs are directly cut by PA-X according to our analysis.

Overall, with 5’ RACE-seq and our updated PyDegradome pipeline (**Fig. 1**), we present a streamlined way to identify and characterize endoRNase cut sites throughout the transcriptome that can be easily adapted to study other RNases. This method can uncover cut site preference characteristics that are masked *in vitro*, because RNase activity *in vitro* can be pushed to the limit by providing sufficient substrate or because cofactors are missing. This method directly identifies cleaved RNAs, instead of relying on RNA steady state levels, which can come from both degradation and transcription repression. Using this method, we have identified a new level of specificity in the mechanism of action of PA-X, adding to our understanding of how PA-X selects RNAs for degradation and distinguishes between host and viral mRNAs. This knowledge brings us one step closer to understanding how PA-X drives host shutoff, and more generally how PA-X modulates inflammation and influenza pathogenesis. It also highlights the importance of determining the cleavage specificity of RNases to understand their functional role, and reveals the potential for viral self vs. non-self discrimination mechanisms.

## Supporting information

Supplemental_figures_and_TablesS1,S3

## Acknowledgments

We thank Albert Tai and the personnel of the Tufts University Core Facility - Genomics Core for help with the sequencing. We thank Drs. Craig McCormick and Denys Khaperskyy and members of their laboratories for their advice and input. We thank Drs. Richard Webby, Gideon Dreyfuss, Jesse Bloom and Seema Lakdawala for constructs, Dr. Mariano Garcia Blanco for providing the Northern Blot protocol, Dr. Bernard Moss for cell lines, and Drs. John Coffin, Claire Moore, Karl Munger and members of the Gaglia lab for critical reading of the manuscript. This work was supported by NIH grant R01 AI137358 (to M.M.G.). L.G. was supported by NIH F31 AI154587 and T32 GM007310. CHR was partially supported by the Applied Mathematics Program of the U.S. DOE Office of Science Advanced Scientific Computing Research under contract number DE-AC02-05CH11231.

## Author contribution

Conceptualization, L.G. and M.M.G.; Methodology, L.G., C.H.R. and M.M.G.; Investigation, L.G., A.I. and I.G.; Writing – Original Draft, L.G. and M.M.G.; Writing – Review & Editing, A.I., I.G. and C.H.R.; Funding Acquisition, M.M.G.; Supervision, M.M.G.

## Declaration of interests

The authors declare no competing interests.

## Online Methods

### Plasmids

pCR3.1-PA-X-myc, pCR3.1-PA-X-D108A-myc and pCR3.1-PA(fs)-myc (all sequences from the PR8 strain) were previously described^50^. pCDEF3-SOX and pCDNA3.1-vhs were previously described^51, 52^. The luciferase constructs with and without the !-globin intron were a kind gift from Dr. Gideon Dreyfuss^26^. Gibson cloning using HiFi assembly mix (New England Biolabs) was used to make all other constructs, unless otherwise stated. PR8 pHW-PA(ΔX) plasmid was generated as previously described^4^ from pHW-193, a kind gift from Dr. R. Webby (St Jude’s Children Research Hospital, Memphis TN). pCR3.1-PA-X-Perth-myc was generated by PCR amplifying pHW-Perth09-PA, a kind gift from Dr. S. Lakdawala^53^ while removing one nucleotide at the frameshift sequence. Fragments were then introduced into the pCR3.1-PA-X-myc vector digested with SalI and MluI to excise PR8 PA-X. pHW-Perth-PA(ΔX) was generated from pHW-Perth09- PA by inserting the ΔX mutations into PCR primers and reintroducing the fragments into pHW- Perth09-PA digested with NheI and BamHI. The same strategy was used to generate pSJ560- TN/CA/7-PA(ΔX) from NheI digested pSJ560-TN/CA/7-PA, a kind gift from Dr. R. Webby (St Jude’s Children Research Hospital, Memphis TN). 99 base pair DNA sequences from genes selected for validation were amplified from human RNA using SuperScript IV One-STEP RT- PCR (Thermo Fisher Scientific), or PCR amplified from human cDNA using Vent Polymerase (New England Biolabs), then introduced into the luciferase with an intron construct at the EcoRI site. The GCTG cut site was mutated to TAGC using the QuickChange site directed mutagenesis kit (Agilent). 51 base pair constructs were generated by PCR amplification of their respective 99 base pair constructs, then inserted into the luciferase EcoRI site by DNA ligation with T4 DNA ligase (New England Biolabs). 27 base pair constructs were generated by annealing primers together and inserting them into the luciferase EcoRI site by DNA ligation with T4 DNA ligase. 15 base pair constructs were generated by introducing the new 15 base pair sequence into the reverse primer and PCR amplifying the rest of the luciferase sequence between the HindIII and EcoRI sites. The hairpin loop structure mutants were generated by inserting targeted mutations into primers and amplifying the cut site sequences with these primers with overlapping sequences to assemble the inserts back to the HindIII/EcoRI digested luciferase vector. The 99 base pair validation sequences were introduced into the !-globin intron of the luciferase construct by PCR amplification of the upstream luciferase sequence from the HindIII site, the 99 base pair sequence, and the downstream luciferase sequence up to the EcoRI site, all with overlapping sequences to assemble the fragments. The WT STOML2 51 bp cut site sequence was introduced into pHW- 197-PR8-M at the end of each coding sequence by PCR amplification of the upstream M sequence from the BamHI site, the STOML2 51 bp sequence, a repeat of the full packaging sequence, and the downstream plasmid sequence up to the BstEII site, all with overlapping sequences to assemble the fragments. The construct containing the mutant STOML2 51 bp cut site in was generated from the WT sequence containing construct by mutating the GCTG sequence at the cut site to TAGC using the QuickChange site directed mutagenesis kit (Agilent). All primers used for cloning are listed in Supplementary Table 3.

### Cell lines and transfections

Human embryonic kidney HEK293T cells and Madin-Darby Canine Kidney (MDCK) cells were commercially obtained (ATCC). Wild type and Xrn1 knock out human adenocarcinoma alveolar basal epithelial (A549) cells were a kind gift from Dr. Bernard Moss^22^. HEK293T and MDCK cells are female and A549 cells are male. All cells were maintained in Dulbecco’s modified Eagle’s medium (DMEM) high glucose (Gibco) supplemented with 10% fetal bovine serum (Hyclone) at 37 °C and 5% CO_2_. HEK293T inducible shXrn1 (ishXrn1) cells were previously described^20^. For northern blotting and 5’ RACE validation experiments, HEK293T ishXrn1 cells were treated with 1 μg/ml doxycycline (Thermo Fisher Scientific) for 3-4 days to induce expression of the shRNA, plated on 6-well or 12-well plates, and transfected with 1000 or 800 ng/ml total DNA (including 62.5 or 50 ng/ml PA-X construct, respectively) using jetPRIME transfection reagent (Polyplus transfection, VWR). Cells were harvested 24 hours after transfection for RNA extraction and purification, or infected overnight as described below and harvested the next day for **Extended Data Fig. 6B**.

### Viruses and infections

Wild-type influenza A/Puerto Rico/8/1934 (H1N1) (PR8), A/Tennessee/1-560/2009 (H1N1) (H1N1pdm09) and A/Perth/16/2009 (H3N2) (Perth) viruses, as well as their mutant recombinant virus counterparts PR8-PA(ΔX), H1N1pdm09-PA(ΔX) and Perth-PA(ΔX), and the mutant recombinant viruses containing the WT or mutated STOML2 51 bp cut site in the M or NA segment were generated using the 8-plasmid reverse genetic system^54^ as previously described^4, 55^. Plasmids were gifts from Drs. Webby, Bloom and Lakdawala. Viral stocks were propagated in MDCK cells and infectious titers determined by plaque assays in MDCK cells using 1.2% Avicel overlays as previously described ^56^. We realized that some of the WT PR8 and PR8-PA(ΔX) virus preparations were contaminated with mycoplasma, but also validated our 5’ RACE-seq results with clean preparations of PR8, H1N1pdm09 and Perth viruses. Influenza infections were performed in DMEM supplemented with 0.5% low endotoxin bovine serum albumin (BSA, Sigma-Aldrich), referred to as infection media. For 5’ RACE, western blotting and qRT-PCR experiments, WT or Xrn1 knock out A549 cells or transfected HEK293T ishXrn1 cells were mock-infected or infected with WT, PA(ΔX) or viruses containing the STOML2 sequences at a MOI of 1 in a low volume of media for 1 hour, then more infection media supplemented with 0.5 µg/ml TPCK-treated trypsin (Sigma-Aldrich) was added and cells were incubated for 15 hours at 37 °C in 5% CO_2_ atmosphere. Cells were then collected for RNA isolation or preparation of lysates for western blotting. For time course experiments, MDCK cells were mock-infected or infected with WT PR8, PR8-PA(ΔX) or viruses containing the STOML2 sequences at a MOI of 0.05 in a low volume of media for 1 hour. The inoculum was then removed, cells were washed with PBS, and infection media supplemented with 0.5 µg/ml TPCK-treated trypsin was added. Supernatant and RNA were collected for the 1h time point, while the rest of the cells were incubated at 37 °C in 5% CO_2_ atmosphere until collection of supernatant and RNA at 8h, 24h and 48h post infection. Viral titers were quantified by 50% tissue culture infectious dose (TCID50). Briefly, 10 µl of undiluted (for 1h and 8h) or of a 1:100 dilution (for 24h and 48h) of the viral supernatant was serially diluted 1:10 in a 96-well plate containing 90 µl of infection media. 25 x 10^3^ MDCK cells were then added to each well. Plates were incubated for 4 days at 37 °C in 5% CO_2_ atmosphere before being scored for cytopathic effects. The method of Reed and Muench^57^ was used to calculate viral titers via the Bloom lab Python script at https://github.com/jbloom/reedmuenchcalculator. To test whether the STOML2 sequence was retained, viral genomic RNA was purified from virus preparations and infected cell supernatant using the QIAamp Viral RNA Mini kit (Qiagen) following manufacturer’s protocol. The RNA was then treated with RNase-Free DNase set (Qiagen), purified and concentrated using the RNA Clean & Concentrator kit (Zymo Research), and reverse transcribed to cDNA using iScript Supermix (Bio-Rad), per manufacturer’s protocol. Taq DNA polymerase (New England Biolabs) was used to amplify the region around the STOML2 sequence. PCR products were visualized on a 2% agarose gel, and DNA from bands were sequenced to confirm that there was no mutation in the STOML2 sequences.

### Protein harvesting and western blotting

Cell lysates were prepared using radioimmunoprecipitation assay (RIPA) buffer (50 mM Tris-HCl, pH 7.4, 150 mM NaCl, 2 mM EDTA, 0.5% sodium deoxycholate, 0.1% SDS, 1% NP-40) supplemented with 50 μg/ml phenylmethylsulfonyl fluoride (PMSF; G-Biosciences) and cOmplete protease cocktail inhibitor (Roche). 20-100 µg of protein was loaded on an SDS-PAGE gel (Bio-Rad) and transferred onto PVDF membranes (EMD Millipore), then blocked with 5% milk in phosphate-buffered saline with 0.1% Tween 20 (PBST). Western blots were performed with mouse anti Xrn1 C-1 antibodies (Santa Cruz Biotechnology #sc-165985, 1:500), rabbit anti influenza A virus PA antibodies (GeneTex #125932, 1:1000), or rabbit anti *b*-tubulin 9F3 antibodies (Cell Signaling Technologies #2128, 1:1000) diluted in 0.5% milk PBTS. Secondary antibodies were purchased from Southern Biotech and used at 1:5,000 dilution. Western blots were imaged with a Syngene G:Box chemi-XX6 system (GeneSys software version 1.7.2.0).

### RNA purification, cDNA generation and qRT-PCR

RNA was extracted and purified using either the RNeasy Plus mini kit (Qiagen) for 5’ RACE-seq experiments, or the Quick-RNA miniprep kit (Zymo Research) for all other experiments, following manufacturer’s protocol. RNA was then treated with Turbo DNase (Life Technologies) and extracted from the DNase reaction by adding phenol chloroform, centrifuging at 12,000 x g for 5 min, collecting the aqueous layer and precipitating RNA in ethanol for 1 hour at -20 °C. RNA was then pelleted, washed with 75% ethanol, then resuspended in RNase-free water. For quantitative real time PCR (qRT-PCR) experiments, resuspended RNA was reverse transcribed to cDNA using iScript Supermix (Bio-Rad), per manufacturer’s protocol. qRT-PCR was performed using iTaq Universal SYBR Green Supermix (Bio-Rad), on the Bio-Rad CFX Connect Real-Time System qPCR and analyzed with Bio-Rad CFX Manager 3.1 or CFX Maestro 2.0 programs. The primers used are listed in Supplementary Table 3. Northern blotting, 5’ RACE and 5’ RACE-seq specific RNA processing are detailed in the next sections.

### Northern blotting

Northern blotting was performed using the NorthernMax kit (Invitrogen) solutions. After DNase treatment and phenol chloroform extraction, 5 µg of DNase-treated RNA was separated on a 1.2 % agarose gel, then transferred by capillary blotting onto a Biodyne B nylon membrane (ThermoFisher Scientific). Biotinylated DNA probes against the 3’ UTR of the luciferase reporters were generated by PCR amplification using Taq polymerase (New England Biolabs) in the presence of Biotin-16-dUTP (Sigma-Aldrich), see primers in Supplementary Table 3. Northern blots were probed with these biotinylated DNA probes, then blocked with Intercept PBS Blocking Buffer (LI-COR Biosciences) supplemented with 1 % SDS. Finally, northern blots were incubated with IRDye 800CW Streptavidin (LI-COR Biosciences, 1:10,000) diluted in Intercept PBS Blocking Buffer supplemented with 1 % SDS, then imaged on a LI-COR Odyssey CLx imaging system (Image Studio software version 5.2).

### 5’ RACE

After DNase treatment and phenol chloroform extraction, the RACE adapter (see Supplementary Table 3) was ligated to 1-5 µg RNA using T4 RNA ligase (Invitrogen) for 1 hour at 37 °C or 2 hours at 25 °C. Ligated RNA was reverse transcribed to cDNA using MMLV RT (Thermo Fisher Scientific) per manufacturer’s protocol. Taq DNA polymerase (New England Biolabs) was used to amplify fragments of interest, if present, in two rounds of nested PCRs using forward primers annealing to the RACE adapter (RACE outer and RACE inner, Supplementary Table 3) and reverse primers annealing within the gene of interest (see Supplementary Table 3). Finally, PCR products were run on a 2% agarose gel containing HydraGreen safe DNA dye (ACTGene) to visualize the amplified fragments and imaged with a Syngene G:Box chemi-XX6 system (GeneSys software version 1.7.2.0). DNA fragments from gel bands at the expected sizes were extracted and sequenced to confirm their identities.

### 5’ RACE-seq library preparation and high-throughput sequencing

Our 5’ RACE-seq library preparation protocol was inspired by previous PARE-Seq protocols with some changes^19, 58, 59^. After DNase treatment and phenol chloroform extraction, 3 µg RNA for each sample were mixed with 6 µl of ERCC ExFold RNA spike-in mixes diluted to 1:100 (Invitrogen). Next, ribosomal RNA was removed from each sample using Ribo-Zero Plus kit (Illumina). The rest of the RNA was ligated to the RACE-seq adapter (see Supplementary Table 3) using T4 RNA ligase (Invitrogen) for 2 hours at 25 °C. The RACE-seq adapter contains a unique molecular identifier (UMI) in order to identify and remove reads that are the result of PCR duplication during the library preparation. Ligated RNA was then reverse transcribed to cDNA with SuperScript III RT (Invitrogen) using random primers of different lengths that also include a known sequence adapter (long and short RT primers, Supplementary Table 3). Finally, reads were amplified in two rounds of PCR (six cycles each) to enrich for ligated fragments and add the Illumina adapters, which include library barcodes for multiplexing (PCR1 and PCR2 primers, Supplementary Table 3). Between the two PCR rounds, DNA fragments of 150-400 bp were selected using SPRIselect Reagent (Beckman Coulter). The quality of each library was evaluated using a Fragment Analyzer (Advanced Analytical Technologies, Inc.) at the Tufts University Core Facility - Genomics Core. High-throughput sequencing was carried out by the Tufts Genomics facility on a HiSeq 2500, obtaining single-end 50 nucleotide reads on three total lanes (Supplementary Table 1). The raw sequences and identified cut sites are deposited on the NCBI GEO database (GSE207253, reviewer token: idatiuqodnyxtev).

### Reads preprocessing and alignment

After labeling each read with its own UMI, cutadapt v3.5^60^ was used to trim Illumina adapters and UMI from each read sequence. Trimmed reads were then aligned to the human genome version GRCh38/hg38, the PR8 genome and the ERCC spike sequences using HISAT2 v2.2.1^61^ with the following specific parameters: retain unique alignments only (-k 1 option), disallow softclipping (--no-softclip option), and select fr-secondstrand library type (--rna-strandness F option). Finally, reads with the same UMI mapping to the same genome region were collapsed into one, to eliminate read duplications that can arise from PCR amplification. Of note, only limited duplication was observed. Uniquely aligned reads were then used for the PyDegradome pipeline (Supplementary Table 1). Reads that aligned to the wrong strand (based on their strandness) and contained insertions or deletions were discarded for the PyDegradome analysis.

### PyDegradome and other python analyses

PyDegradome is a Python-based peak finding pipeline developed to identify ribonuclease cut sites, which we originally reported^20^ and modified here. Once all reads are aligned to the genome, the position of the first nucleotide of every read is recorded, and the number of reads that map to the same position is counted for each read of each library. PyDegradome then uses a Bayesian probability model to identify which positions have statistically significantly more reads mapping to them in a test sample *vs.* a control sample. This significance can be made more or less stringent based on three parameters that can be optimized by the user: multiplicative factor, confidence level and scanning window. The multiplicative factor determines by how much the read count at a specific nucleotide position in the test sample needs to exceed the count in the control sample. The confidence level sets the cutoff of statistical significance. The multiplicative factor and confidence level together define the threshold that the test sample needs to exceed to be considered significantly above background. The scanning window determines how many consecutive nucleotides need to have an average read count above the threshold value for a peak to be called. This parameter removes isolated high read counts that could skew the analysis. All the data showed here use a window of 4 nucleotides. For a more detailed explanation of PyDegradome, see our previous PyDegradome publication^20^. In this publication, we updated our older version of PyDegradome to be usable with the alignment tool HISAT2 and the human genome GRCh38/hg38. Our older version of PyDegradome also restricted the cut site analysis to exons, and we have updated it to include introns in case PA-X cut pre-mRNA. The PyDegradome 2.0 updated script can be found in our laboratory’s GitHub page (https://github.com/mgaglia81/PyDegradome). The down-stream motif generation and scoring analysis was not modified and is still available to download as supplementary material in our previous article^20^ or our GitHub page. To calculate the percentage of GCUG within transcriptomes, we counted the number of GCUG tetramers found in each RNA sequences and divided this number by the total possible number of tetramers in that RNA, i.e. length of RNA minus 3. To calculate the GCTG percentage in genomes, we counted the number of GCTG / GCUG tetramers in each chromosome / influenza segment and divided this number by the total number of tetramers in that chromosome / influenza segment, i.e. length of chromosome / segment minus 3. The python script used is available on our GitHub page. Transcriptome fasta files were downloaded from the Ensembl^62^ ftp server available at https://ftp.ensembl.org/pub/current_fasta/. Swan goose: *Anser cygnoides* GooseV1.0, INSDC Assembly GCA_002166845.1, Jun 2017 by Poultry Science Institute. Pig: *Sus scrofa* Sscrofa11.1, INSDC Assembly GCA_000003025.6, Feb 2017 by the Swine Genome Sequencing Consortium (SGSC)^63, 64^. Chicken: *Gallus gallus* bGalGal1.mat.broiler.GRCg7b, INSDC Assembly GCA_016699485.1, Jan 2021 by Vertebrate Genomes Project. Mallard duck: *Anas platyrhynchos* ASM874695v1, INSDC Assembly GCA_008746955.1, Sep 2019 by China Agricultural University. Influenza transcriptomes were downloaded from the NIAID Influenza Research Database (IRD)^65^ (previously available at http://www.fludb.org). A/Puerto-Rico/8/34 (H1N1) Genbank CY033577-84, A/Tennessee/05/2009 (H1N1) Genbank GQ160528-31 GQ396729-30 GQ894895 GQ200205, A/Perth/16/2009 (H3N2) Genbank KJ609203-10, A/swine/Guangdong/2722/2011 (H1N1) Genbank KM027548-56^66^, A/chicken/Pakistan/UDL-01/2008 (H9N2) Genbank CY038455-62^67^, A/tree sparrow/Jiangsu/1/2008 (H5N1) Genbank GQ202207-14^68^, A/anhui/1/2005 (H5N1) HPAI Genbank HM172438, HM172394, HM172342, HM172104, HM172254, HM172189, HM172159, HM172266^69^.

## Data availability

All primary data except for 5’RACE-seq data are publicly available as a Figshare collection, https://doi.org/10.6084/m9.figshare.c.6392286. As stated above, 5’RACE-seq data is deposited in NCBI GEO database, (GSE207253, reviewer token: idatiuqodnyxtev) and all python scripts used, including the PyDegradome 2.0 code can be found in our laboratory’s GitHub page (https://github.com/mgaglia81/PyDegradome).

## Notes

### Competing Interest Statement

The authors have declared no competing interest.

### Summary of Updates

Additional experiments and analyses; revised figures and text

## References

1. Gaucherand, L. & Gaglia, M. M. The Role of Viral RNA Degrading Factors in Shutoff of Host Gene Expression. Annu Rev Virol (2022) doi:10.1146/annurev-virology-100120-012345.

2. Bercovich-Kinori, A. et al. A systematic view on influenza induced host shutoff. eLife 5, e18311 (2016).

3. Jagger, B. W. et al. An overlapping protein-coding region in influenza A virus segment 3 modulates the host response. Science (2012) doi:10.1126/science.1222213.

4. Gaucherand, L. et al. The Influenza A Virus Endoribonuclease PA-X Usurps Host mRNA Processing Machinery to Limit Host Gene Expression. Cell Reports 27, 776–792.e7 (2019).

5. Gao, H. et al. The contribution of PA-X to the virulence of pandemic 2009 H1N1 and highly pathogenic H5N1 avian influenza viruses. Sci Rep 5, 8262 (2015).

6. Gong, X. Q. et al. PA-X protein decreases replication and pathogenicity of swine influenza virus in cultured cells and mouse models. Veterinary Microbiology 205, 66–70 (2017).

7. Hayashi, T., MacDonald, L. A. & Takimoto, T. Influenza A Virus Protein PA-X Contributes to Viral Growth and Suppression of the Host Antiviral and Immune Responses. Journal of Virology (2015) doi:10.1128/JVI.00319-15.

8. Hu, J. et al. PA-X Decreases the Pathogenicity of Highly Pathogenic H5N1 Influenza A Virus in Avian Species by Inhibiting Virus Replication and Host Response. Journal of Virology (2015) doi:10.1128/JVI.02132-14.

9. Firth, A. E. et al. Ribosomal frameshifting used in influenza A virus expression occurs within the sequence UCC-UUU-CGU and is in the +1 direction. Open Biology 2, 1–7 (2012).

10. Dias, A. et al. The cap-snatching endonuclease of influenza virus polymerase resides in the PA subunit. Nature 458, 914–918 (2009).

11. Yuan, P. et al. Crystal structure of an avian influenza polymerase PA_N_ reveals an endonuclease active site. Nature 458, 909–913 (2009).

12. Bavagnoli, L. et al. The novel influenza A virus protein PA-X and its naturally deleted variant show different enzymatic properties in comparison to the viral endonuclease PA. Nucleic Acids Research 43, 9405–9417 (2015).

13. Khaperskyy, D. A., Schmaling, S., Larkins-Ford, J., McCormick, C. & Gaglia, M. M. Selective Degradation of Host RNA Polymerase II Transcripts by Influenza A Virus PA-X Host Shutoff Protein. PLoS Pathogens 12, (2016).

14. Gaglia, M. M., Covarrubias, S., Wong, W. & Glaunsinger, B. A. A Common Strategy for Host RNA Degradation by Divergent Viruses. Journal of Virology 86, 9527–9530 (2012).

15. Chaimayo, C., Dunagan, M., Hayashi, T., Santoso, N. & Takimoto, T. Specificity and functional interplay between influenza virus PA-X and NS1 shutoff activity. PLOS Pathogens 14, e1007465 (2018).

16. Saïda, F., Uzan, M. & Bontems, F. The phage T4 restriction endoribonuclease RegB: a cyclizing enzyme that requires two histidines to be fully active. Nucleic Acids Research 31, 2751–2758 (2003).

17. Lebars, I., Hu, R.-M., Lallemand, J.-Y., Uzan, M. & Bontems, F. Role of the Substrate Conformation and of the S1 Protein in the Cleavage Efficiency of the T4 Endoribonuclease RegB*. Journal of Biological Chemistry 276, 13264–13272 (2001).

18. Endo, Y., Glück, A., Chan, Y. L., Tsurugi, K. & Wool, I. G. RNA-protein interaction. An analysis with RNA oligonucleotides of the recognition by alpha-sarcin of a ribosomal domain critical for function. J Biol Chem 265, 2216–2222 (1990).

19. Chao, Y. et al. In Vivo Cleavage Map Illuminates the Central Role of RNase E in Coding and Non-coding RNA Pathways. Molecular Cell 65, 39–51 (2017).

20. Gaglia, M. M., Rycroft, C. H. & Glaunsinger, B. A. Transcriptome-Wide Cleavage Site Mapping on Cellular mRNAs Reveals Features Underlying Sequence-Specific Cleavage by the Viral Ribonuclease SOX. PLoS Pathogens 11, 1–25 (2015).

21. Abernathy, E., Gilbertson, S., Alla, R. & Glaunsinger, B. Viral nucleases induce an mRNA degradation-transcription feedback loop in mammalian cells. Cell Host and Microbe 18, 243–253 (2015).

22. Liu, R. & Moss, B. Opposing Roles of Double-Stranded RNA Effector Pathways and Viral Defense Proteins Revealed with CRISPR-Cas9 Knockout Cell Lines and Vaccinia Virus Mutants. Journal of Virology 90, 7864–7879 (2016).

23. Hayashi, T., Chaimayo, C., McGuinness, J. & Takimoto, T. Critical Role of the PA-X C- Terminal Domain of Influenza A Virus in Its Subcellular Localization and Shutoff Activity. Journal of Virology (2016) doi:10.1128/JVI.00954-16.

24. Gao, H. et al. PA-X is a virulence factor in avian H9N2 influenza virus. J. Gen. Virol. 96, 2587–2594 (2015).

25. Rigby, R. E., Wise, H. M., Smith, N., Digard, P. & Rehwinkel, J. PA-X antagonises MAVS- dependent accumulation of early type I interferon messenger RNAs during influenza A virus infection. Sci Rep 9, 7216 (2019).

26. Younis, I. et al. Rapid-response splicing reporter screens identify differential regulators of constitutive and alternative splicing. Molecular and cellular biology 30, 1718–28 (2010).

27. Levene, R. E., Shrestha, S. D. & Gaglia, M. M. The influenza A virus host shutoff factor PA- X is rapidly turned over in a strain-specific manner. J Virol 95, e02312–20 (2021).

28. Huang, L. et al. LinearFold: linear-time approximate RNA folding by 5’-to-3’ dynamic programming and beam search. Bioinformatics 35, i295–i304 (2019).

29. Do, C. B., Woods, D. A. & Batzoglou, S. CONTRAfold: RNA secondary structure prediction without physics-based models. Bioinformatics 22, e90–e98 (2006).

30. Lorenz, R. et al. ViennaRNA Package 2.0. Algorithms for Molecular Biology 6, 1–14 (2011).

31. Mathews, D. H. et al. Incorporating chemical modification constraints into a dynamic programming algorithm for prediction of RNA secondary structure. Proceedings of the National Academy of Sciences of the United States of America 101, 7287 (2004).

32. Plotch, S. J., Bouloy, M., Ulmanen, I. & Krug, R. M. A unique cap(m7GpppXm)-dependent influenza virion endonuclease cleaves capped RNAs to generate the primers that initiate viral RNA transcription. Cell 23, 847–858 (1981).

33. Datta, K., Wolkerstorfer, A., Szolar, O. H. J., Cusack, S. & Klumpp, K. Characterization of PA-N terminal domain of Influenza A polymerase reveals sequence specific RNA cleavage. Nucleic Acids Research 41, 8289–8299 (2013).

34. Sikora, D., Rocheleau, L., Brown, E. G. & Pelchat, M. Deep sequencing reveals the eight facets of the influenza A/HongKong/1/1968 (H3N2) virus cap-snatching process. Scientific Reports 4, (2014).

35. Sikora, D., Rocheleau, L., Brown, E. G. & Pelchat, M. Influenza A virus cap-snatches host RNAs based on their abundance early after infection. Virology 509, 167–177 (2017).

36. Gu, W. et al. Influenza A virus preferentially snatches noncoding RNA caps. RNA 21, 2067– 2075 (2015).

37. Brannan, K. et al. MRNA Decapping Factors and the Exonuclease Xrn2 Function in Widespread Premature Termination of RNA Polymerase II Transcription. Molecular Cell 46, 311–324 (2012).

38. Wang, X.-H. et al. The Role of PA-X C-Terminal 20 Residues of Classical Swine Influenza Virus in Its Replication and Pathogenicity. Veterinary Microbiology 251, 108916 (2020).

39. Oishi, K., Yamayoshi, S. & Kawaoka, Y. Identification of amino acid residues in influenza A virus PA-X that contribute to enhanced shutoff activity. Frontiers in Microbiology 10, 1–8 (2019).

40. Feng, K. H., Sun, M., Iketani, S., Holmes, E. C. & Parrish, C. R. Comparing the functions of equine and canine influenza H3N8 virus PA-X proteins: Suppression of reporter gene expression and modulation of global host gene expression. Virology (2016) doi:10.1016/j.virol.2016.06.001.

41. Nogales, A. et al. Natural Selection of H5N1 Avian Influenza A Viruses with Increased PA- X and NS1 Shutoff Activity. Viruses 2021, Vol. 13, Page 1760 13, 1760 (2021).

42. Wilamowski, M., Gorecki, A., Dziedzicka-Wasylewska, M. & Jura, J. Substrate specificity of human MCPIP1 endoribonuclease. Sci Rep 8, 7381 (2018).

43. Lee, N. et al. Genome-wide analysis of influenza viral RNA and nucleoprotein association. Nucleic Acids Res 45, 8968–8977 (2017).

44. Takata, M. A. et al. CG dinucleotide suppression enables antiviral defence targeting non-self RNA. Nature 550, 124–127 (2017).

45. Pezda, A. C., Penn, A., Barton, G. M. & Coscoy, L. Suppression of TLR9 immunostimulatory motifs in the genome of a gammaherpesvirus. J Immunol 187, 887–896 (2011).

46. Hartenian, E. & Glaunsinger, B. A. Feedback to the central dogma: cytoplasmic mRNA decay and transcription are interdependent processes. https://doi.org/10.1080/10409238.2019.167908354, 385–398 (2019).

47. Duncan-Lewis, C., Hartenian, E., King, V. & Glaunsinger, B. A. Cytoplasmic mrna decay represses rna polymerase ii transcription during early apoptosis. eLife 10, (2021).

48. Zhao, N. et al. Influenza virus infection causes global RNAPII termination defects. Nature Structural and Molecular Biology 25, 885–893 (2018).

49. Bauer, D. L. V. et al. Influenza Virus Mounts a Two-Pronged Attack on Host RNA Polymerase II Transcription. Cell Reports 23, 2119–2129.e3 (2018).

50. Khaperskyy, D. A. et al. Influenza A Virus Host Shutoff Disables Antiviral Stress-Induced Translation Arrest. PLOS Pathogens 10, e1004217 (2014).

51. Glaunsinger, B. & Ganem, D. Lytic KSHV infection inhibits host gene expression by accelerating global mRNA turnover. Mol. Cell 13, 713–723 (2004).

52. Jones, F. E., Smibert, C. A. & Smiley, J. R. Mutational analysis of the herpes simplex virus virion host shutoff protein: evidence that vhs functions in the absence of other viral proteins. J Virol 69, 4863–4871 (1995).

53. Le Sage, V. et al. Cell-Culture Adaptation of H3N2 Influenza Virus Impacts Acid Stability and Reduces Airborne Transmission in Ferret Model. Viruses 13, 719 (2021).

54. Hoffmann, E., Neumann, G., Kawaoka, Y., Hobom, G. & Webster, R. G. A DNA transfection system for generation of influenza A virus from eight plasmids. Proceedings of the National Academy of Sciences 97, 6108–6113 (2000).

55. Khaperskyy, D. A., Hatchette, T. F. & McCormick, C. Influenza A virus inhibits cytoplasmic stress granule formation. The FASEB Journal 26, 1629–1639 (2012).

56. Matrosovich, M., Matrosovich, T., Garten, W. & Klenk, H. D. New low-viscosity overlay medium for viral plaque assays. Virology Journal 3, 1–7 (2006).

57. Reed, L. J. & Muench, H. A SIMPLE METHOD OF ESTIMATING FIFTY PER CENT ENDPOINTS12. American Journal of Epidemiology 27, 493–497 (1938).

58. German, M. A., Luo, S., Schroth, G., Meyers, B. C. & Green, P. J. Construction of Parallel Analysis of RNA Ends (PARE) libraries for the study of cleaved miRNA targets and the RNA degradome. Nat Protoc 4, 356–362 (2009).

59. Zhai, J., Arikit, S., Simon, S. A., Kingham, B. F. & Meyers, B. C. Rapid construction of parallel analysis of RNA end (PARE) libraries for Illumina sequencing. Methods 67, 84–90 (2014).

60. Martin, M. Cutadapt removes adapter sequences from high-throughput sequencing reads. EMBnet.journal 17, 10–12 (2011).

61. Kim, D., Paggi, J. M., Park, C., Bennett, C. & Salzberg, S. L. Graph-based genome alignment and genotyping with HISAT2 and HISAT-genotype. Nature Biotechnology 37, 907–915 (2019).

62. Cunningham, F., et al. Ensembl 2022. Nucleic Acids Research 50, D988–D995 (2022).

63. Warr, A. et al. An improved pig reference genome sequence to enable pig genetics and genomics research. 668921 Preprint at https://doi.org/10.1101/668921 (2019).

64. Li, M. et al. Comprehensive variation discovery and recovery of missing sequence in the pig genome using multiple de novo assemblies. Genome Res. 27, 865–874 (2017).

65. Zhang, Y. et al. Influenza Research Database: An integrated bioinformatics resource for influenza virus research. Nucleic Acids Res 45, D466–D474 (2017).

66. Liang, H. et al. Expansion of genotypic diversity and establishment of 2009 H1N1 pandemic-origin internal genes in pigs in China. J Virol 88, 10864–10874 (2014).

67. Iqbal, M., Yaqub, T., Reddy, K. & McCauley, J. W. Novel genotypes of H9N2 influenza A viruses isolated from poultry in Pakistan containing NS genes similar to highly pathogenic H7N3 and H5N1 viruses. PLoS One 4, e5788 (2009).

68. Liu, Q. et al. Characterization of a highly pathogenic avian influenza H5N1 clade 2.3.4 virus isolated from a tree sparrow. Virus Res 147, 25–29 (2010).

69. Li, Y. et al. Continued evolution of H5N1 influenza viruses in wild birds, domestic poultry, and humans in China from 2004 to 2009. J Virol 84, 8389–8397 (2010).

70. Crooks, G. E., Hon, G., Chandonia, J. M. & Brenner, S. E. WebLogo: A Sequence Logo Generator. Genome Research 14, 1188–1190 (2004).

