## Supplemental_figures_and_TablesS1,S3 for "The cut site specificity of the influenza A virus endoribonuclease PA-X allows it to discriminate between host and viral mRNAs"

Supplementary Table 1 – Summary of RACE-seq read data.

Number of total reads sequenced, reads that aligned exactly one time to hg38/ERCC spike ins/PR8, and reads used in PyDegradome analysis for each sample.

| Replicate | Sample | Total | Uniquely aligned | % uniquely aligned | Used as PyDegradome input |
|---|---|---|---|---|---|
| 1 | Mock | 49956233 | 29851634 | 59.8% | 26375238 |
| 1 | WT PR8 | 59832882 | 30563534 | 51.1% | 18055280 |
| 1 | PR8-PA(ΔX) | 61091210 | 32019504 | 52.4% | 22210910 |
| 2 | Mock | 46603110 | 27547835 | 59.1% | 23708011 |
| 2 | WT PR8 | 50920984 | 28938148 | 56.8% | 16459354 |
| 2 | PR8-PA(ΔX) | 62507002 | 32229277 | 51.6% | 21897094 |
| 3 | Mock | 57788531 | 37176976 | 64.3% | 31313003 |
| 3 | WT PR8 | 60000926 | 32354246 | 53.9% | 17821338 |
| 3 | PR8-PA(ΔX) | 53011926 | 27782595 | 52.4% | 18229661 |

Supplementary Table 2 – Summary of PA-X cut sites (excel sheet)

Supplementary Table 3 – Primers and RNA adapters.

| Name | Sequence (5' → 3') | Source |
|---|---|---|
| RACE-seq RNA adapter | CCCUACACGACGCUCUUCCGAUCUNNNNNNNNN | This paper |
| Short RT primer | TTCAGACGTGTGCTCTTCCGATCNNNNNNNN | This paper |
| Long RT primer | TTCAGACGTGTGCTCTTCCGATCNNNNNNNNNNNNNN | This paper |
| PCR1 F primer | CCCTACACGACGCTCTTCCGATCT | This paper |
| PCR1 R primer | TTCAGACGTGTGCTCTTCCGATC | This paper |
| PCR2 F primer | AATGATACGGCGACCACCGAGATCTACACTCTTTCCCT<br>ACACGACGCTCTTCCGATCT | This paper |
| PCR2 R1 primer (mock rep. 1) | CAAGCAGAAGACGGCATACGAGATTCAAGTGTGACTG<br>GAGTTCAGACGTGTGCTCTTCCGATC | This paper |
| PCR2 R2 primer (WT rep. 1) | CAAGCAGAAGACGGCATACGAGATCTGATCGTGACTG<br>GAGTTCAGACGTGTGCTCTTCCGATC | This paper |
| PCR2 R3 primer (ΔX rep. 1) | CAAGCAGAAGACGGCATACGAGATAAGCTAGTGACTG<br>GAGTTCAGACGTGTGCTCTTCCGATC | This paper |
| PCR2 R4 primer (mock rep. 2) | CAAGCAGAAGACGGCATACGAGATCGTGATGTGACTG<br>GAGTTCAGACGTGTGCTCTTCCGATC | This paper |
| PCR2 R5 primer (WT rep. 2) | CAAGCAGAAGACGGCATACGAGATACATCGGTGACTG<br>GAGTTCAGACGTGTGCTCTTCCGATC | This paper |
| PCR2 R6 primer (ΔX rep. 2) | CAAGCAGAAGACGGCATACGAGATGCCTAAGTGACTG<br>GAGTTCAGACGTGTGCTCTTCCGATC | This paper |
| PCR2 R7 primer (mock rep. 3) | CAAGCAGAAGACGGCATACGAGATTGGTCAGTGACTG<br>GAGTTCAGACGTGTGCTCTTCCGATC | This paper |
| PCR2 R8 primer (WT rep. 3) | CAAGCAGAAGACGGCATACGAGATCACTGTGTGACTG<br>GAGTTCAGACGTGTGCTCTTCCGATC | This paper |
| PCR2 R9 primer (ΔX rep. 3) | CAAGCAGAAGACGGCATACGAGATATTGGCGTGACTG<br>GAGTTCAGACGTGTGCTCTTCCGATC | This paper |
| RACE RNA adapter | GCUGAUGGCGAUGAAUGAACACUGCGUUUGCUGGCU<br>UUGAUGAAA | This paper |
| BCAP31 outer R primer | CTTTGCGTGCTCCTCCAGCAAG | This paper |
| BCAP31 inner R primer | CCGCCACTGTGCTGGATATCTGCAGAATTCCATGGCC<br>AGAACCTGGTTTTC | This paper |
| INSIG1 outer R primer | CAAATTCCATAGGGCTGTGG | This paper |

| | | |
|---|---|---|
| INSIG1 inner R primer | CCGCCACTGTGCTGGATATCTGCAGAATTCCACATGG CAGACAGGTATTCA | This paper |
| SLC7A5 outer R primer | TCCATCCTCCATAGGCAAAGAG | This paper |
| SLC7A5 inner R primer | CCGCCACTGTGCTGGATATCTGCAGAATTCCCACATC CAGTTTGGTGCCTTC | This paper |
| STOML2 outer R primer | GCATCTGCATAGACTCTTTCACC | This paper |
| STOML2 inner R primer | CCGCCACTGTGCTGGATATCTGCAGAATTCCTTGATCT CATAACGGAGGCAG | This paper |
| TUBA1B outer R primer | ATGCTGCAGGGCCAAAAGGAATG | This paper |
| TUBA1B inner R primer | CCGCCACTGTGCTGGATATCTGCAGAATTCCCAACAG AATCCACACCAACCTC | This paper |
| YKT6 outer R primer | CCATAGGTGGGGTGCTGGGAATC | This paper |
| YKT6 inner R primer | CCGCCACTGTGCTGGATATCTGCAGAATTCCTTGTTCA GATGCATAGGCCTC | This paper |
| Luciferase outer R primer | CCGCCACTGTGCTGGATATCTGCAGAATTCGCGCAAG CAGCAGGGTGTCTATCC | This paper |
| Luciferase inner R primer | CCGCCACTGTGCTGGATATCTGCAGAATTCGGGCAGG GTGCCGGCGGCCTG | This paper |
| Luciferase outer int R primer | GTCGAAGATGTTGGGGTGTTG | This paper |
| Luciferase inner int R primer | CCGCCACTGTGCTGGATATCTGCAGAATTCCGTGTTG CAGCAGGATGCTCTC | This paper |
| PR8-HA-GCTG outer R primer | GTCGCATTCTGGGTTTCCCAAG | This paper |
| PR8-HA-GCTG inner R primer | CCGCCACTGTGCTGGATATCTGCAGAATTCCCCCAATT GTAGTGGGGCTATTC | This paper |
| PR8-NS-GCTG outer R primer | GTTCCCGCCATTTCTCGTTTCTG | This paper |
| PR8-NS-GCTG inner R primer | CCGCCACTGTGCTGGATATCTGCAGAATTCCCTCATTA CTGCTTCTCCAAGC | This paper |
| H1N1pdm09-NP-GCTG outer R primer | GATCCATTCCGGTGCGAACAAG | This paper |
| H1N1pdm09-NP-GCTG inner R primer | CCGCCACTGTGCTGGATATCTGCAGAATTCCCTGTAA GACCTGCTGTTGCATCTTCG | This paper |
| H1N1pdm09-PA-GCTG outer R primer | GTGCATGTGTGAGGAAGGAGTTG | This paper |
| H1N1pdm09-PA-GCTG inner R primer | CCGCCACTGTGCTGGATATCTGCAGAATTCCCTCCAG GGATCATTAATCAGGCACTC | This paper |
| PR8-NA STOML2 cut site outer R primer | GTATCACTATTCACGCCAC | This paper |
| PR8-NA STOML2 cut site inner R primer | CCGCCACTGTGCTGGATATCTGCAGAATTCCGAAATG CTGCTCGCACTAGTC | This paper |
| Luciferase northern blot probe F primer | GCTTGCAAGAACTGGTTCAGTAGC | This paper |
| Luciferase northern blot probe R primer | CTTATCATGTCTGCTCGAAGCGG | This paper |
| G6PD qPCR F primer | TGAGCCAGATAGGCTGGAA | Hu BMC Cancer 2013[4] |
| G6PD qPCR R primer | TAACGCAGGCGATGTTGTC | Hu BMC Cancer 2013[4] |
| Luciferase qPCR F primer | ATCGAGGTGGACATTACCTACG | Khaperskyy et al. *PPath* 2016[5] |
| Luciferase qPCR R primer | CGCTCGTTGTAGATGTCGTTAG | Khaperskyy et al. *PPath* 2016[5] |
| Luciferase int qPCR F primer | CGTGGACCGGCTGAAGAGCCTG | This paper |
| Luciferase int qPCR R primer | GTCAGTAAGACCAATAGGTGC | This paper |
| PR8-HA qPCR F primer | CTGGACCTTGCTAAAACCCG | Slaine et al. *JVI* 2021[6] |
| PR8-HA qPCR R primer | TCTGGAAAGGGAGACTGCTG | Slaine et al. *JVI* 2021[6] |
| PR8-NA qPCR F primer | TCACTTGGAATGCAGGACCT | Slaine et al. *JVI* 2021[6] |
| PR8-NA qPCR R primer | CGATTGTTAGCCAGCCCATG | Slaine et al. *JVI* 2021[6] |

| | | |
|---|---|---|
| PR8-M1 qPCR F primer | TTTGGCCTGGTATGTGCAAC | Khaperskyy et al. *PPath* 2016[5] |
| PR8-M1 qPCR R primer | ACCATTTGCCTAGCCTGACT | Khaperskyy et al. *PPath* 2016[5] |
| PR8-M2 qPCR F primer | GGTCGAAACGCCTATCAGAA | Khaperskyy et al. *PPath* 2016[5] |
| PR8-M2 qPCR R primer | ACTTTGGCACTCCTTCCGTA | Khaperskyy et al. *PPath* 2016[5] |
| Luciferase northern blot probe F primer | GCTTGCAAGAACTGGTTCAGTAGC | This paper |
| Luciferase northern blot probe R primer | CTTATCATGTCTGCTCGAAGCGG | This paper |
| Luciferase + STOML2 99 bp cloning PCR1 F | TAGGGAGACCCAAGCTGGCTAGTTAAGCTTGGCAATCCGGTACTGTTGGT | This paper |
| Luciferase + STOML2 99 bp cloning PCR1 R | TCCTTGAGACTCTGCACATAGAATTCACGGCGATCTTGCC | This paper |
| Luciferase + STOML2 99 bp cloning PCR2 F | GGCAAGATCGCCGTGAATTCTATGTGCAGAGTCTCAAGGA | This paper |
| Luciferase + STOML2 99 bp cloning PCR2 R | CTGAACCAGTTCTTGCAAGCAGAATTCCAGGTAAAGGACTCCATCGA | This paper |
| Luciferase + YKT6 99 bp cloning PCR1 F | Same as Luciferase + STOML2 99 bp cloning PCR1 F | This paper |
| Luciferase + YKT6 99 bp cloning PCR1 R | CATCCCCCTTTCCTGGGCTCAGTTGAATTCACGGCGATCTTGCC | This paper |
| Luciferase + YKT6 99 bp cloning PCR2 F | GGCAAGATCGCCGTGAATTCAACTGAGCCCAGGAAAGGGGGATG | This paper |
| Luciferase + YKT6 99 bp cloning PCR2 R | CAGTTCTTGCAAGCAGAATTCGCAGCTGAGTGCACCTGTGTTTG | This paper |
| Luciferase + BCAP31 99 bp cloning PCR1 F | Same as Luciferase + STOML2 99 bp cloning PCR1 F | This paper |
| Luciferase + BCAP31 99 bp cloning PCR1 R | CAGGCGTCTAAGCAGGAAGGAGAATTCACGGCGATCTTGCC | This paper |
| Luciferase + BCAP31 99 bp cloning PCR2 F | GGCAAGATCGCCGTGAATTCTCCTTCCTGCTTAGACGCCTG | This paper |
| Luciferase + BCAP31 99 bp cloning PCR2 R | CTGAACCAGTTCTTGCAAGCAGAATTCCTCACTAGCACTCTCCGCCTG | This paper |
| Luciferase + TUBA1B 99 bp cloning PCR1 F | Same as Luciferase + STOML2 99 bp cloning PCR1 F | This paper |
| Luciferase + TUBA1B 99 bp cloning PCR1 R | CCAGGTCTCCACCAGGCACCGAATTCACGGCGATCTTGCCGCCCTTCTTG | This paper |
| Luciferase + TUBA1B 99 bp cloning PCR2 F | CAAGAAGGGCGGCAAGATCGCCGTGAATTCGGTGCCTGGTGGAGACCTGGCCAA | This paper |
| Luciferase + TUBA1B 99 bp cloning PCR2 R | CTGAACCAGTTCTTGCAAGCAGAATTCAACTTGTGGTCCAGGCGAGC | This paper |
| Luciferase + INSIG1 99 bp cloning PCR1 F | Same as Luciferase + STOML2 99 bp cloning PCR1 F | This paper |
| Luciferase + INSIG1 99 bp cloning PCR1 R | GTGACCTCTCTATAATCACTGAATTCACGGCGATCTTGCCG | This paper |
| Luciferase + INSIG1 99 bp cloning PCR2 F | GCAAGATCGCCGTGAATTCAGTGATTATAGAGAGGTCACAC | This paper |
| Luciferase + INSIG1 99 bp cloning PCR2 R | CAGTTCTTGCAAGCAGAATTCTGTGACCATGTAAACACGCCAC | This paper |
| Luciferase + SLC7A5 99 bp cloning PCR1 F | Same as Luciferase + STOML2 99 bp cloning PCR1 F | This paper |
| Luciferase + SLC7A5 99 bp cloning PCR1 R | GAAGAGCGGCTTGAGCGAATTCACGGCGATCTTGCCG | This paper |
| Luciferase + SLC7A5 99 bp cloning PCR2 F | CAAGATCGCCGTGAATTCGCTCAAGCCGCTCTTCCCCAC | This paper |
| Luciferase + SLC7A5 99 bp cloning PCR2 R | CAGTTCTTGCAAGCAGAATTCCAACCCCCGGCCCGCGCCCCGTACT | This paper |
| Luciferase + STOML2-mut 99 bp cloning F | CATCAACGTGCCTGAGCAGTCGTAGCTGACTCTCGACAATGTAACTCT | This paper |
| Luciferase + STOML2-mut 99 bp cloning R | AGAGTTACATTGTCGAGAGTCAGCTACGACTGCTCAGGCACGTTGATG | This paper |

| | | |
|---|---|---|
| Luciferase + YKT6-mut 99 bp cloning F | GGGATGTTTTCCTGGTGTTTGGATGGTCATAGCGGAGTGTCCATCATC | This paper |
| Luciferase + YKT6-mut 99 bp cloning R | GATGATGGACACTCCGCTATGACCATCCAAACACCAGGAAAACATCCC | This paper |
| Luciferase + BCAP31-mut 99 bp cloning F | CATTTCGCAGCAGGCCACTAGCCTGGCCTCCAATGAAGCC | This paper |
| Luciferase + BCAP31-mut 99 bp cloning R | GGCTTCATTGGAGGCCAGGCTAGTGGCCTGCTGCGAAATG | This paper |
| Luciferase + TUBA1B-mut 99 bp cloning F | GCCAAGGTACAGAGAGCTGTGTGCATTAGCAGCAACACCACAGC | This paper |
| Luciferase + TUBA1B-mut 99 bp cloning R | GCTGTGGTGTTGCTGCTAATGCACACAGCTCTCTGTACCTTGGC | This paper |
| Luciferase + STOML2 51 bp cloning PCR1 F | Same as Luciferase + STOML2 99 bp cloning PCR1 F | This paper |
| Luciferase + STOML2 51 bp cloning PCR1 R | GAATTCACGGCGATCTTGCC | This paper |
| Luciferase + STOML2 51 bp cloning PCR2 F | GGCAAGATCGCCGTGAATTCGTCATCAACGTGCCTGAG | This paper |
| Luciferase + STOML2 51 bp cloning PCR2 R | CAGTTCTTGCAAGCAGAATTCCAGAGTTACATTGTCGAG | This paper |
| Luciferase + YKT6 51 bp cloning PCR1 F | Same as Luciferase + STOML2 99 bp cloning PCR1 F | This paper |
| Luciferase + YKT6 51 bp cloning PCR1 R | Same as Luciferase + STOML2 51 bp cloning PCR1 R | This paper |
| Luciferase + YKT6 51 bp cloning PCR2 F | GGCAAGATCGCCGTGAATTCTTTTCCTGGTGTTTGGATG | This paper |
| Luciferase + YKT6 51 bp cloning PCR2 R | CAGTTCTTGCAAGCAGAATTCTCTTCCCCTGATGATGGAC | This paper |
| Luciferase + BCAP31 51 bp cloning PCR F | GCGGCAAGATCGCCGTGAATTCACTCTCATTTCGCAGCAGGCCA | This paper |
| Luciferase + BCAP31 51 bp cloning PCR R | ACCAGTTCTTGCAAGCAGAATTCTTTAAAGGCTTCATTGGAGGC | This paper |
| Luciferase + SLC7A5 51 bp cloning PCR F | GGCAAGATCGCCGTGAATTCCCCGGTGCCCGAGGAGGCA | This paper |
| Luciferase + SLC7A5 51 bp cloning PCR R | ACCAGTTCTTGCAAGCAGAATTCAGCAGCACGCAGAGGCAGGC | This paper |
| Luciferase + STOML2 27 bp cloning PCR F | GGCAAGATCGCCGTGAATTCCCTGAGCAGTCGGCTGTGACTCTC | This paper |
| Luciferase + STOML2 27 bp cloning PCR R | CAGTTCTTGCAAGCAGAATTCGTCGAGAGTCACAGCCGACTGCTC | This paper |
| Luciferase + YKT6 27 bp cloning PCR F | GGCAAGATCGCCGTGAATTCTTGGATGGTCAGCTGGGAGTGTCC | This paper |
| Luciferase + YKT6 27 bp cloning PCR R | CAGTTCTTGCAAGCAGAATTCGATGGACACTCCCAGCTGACCATC | This paper |
| Luciferase + BCAP31 27 bp cloning PCR F | GGCAAGATCGCCGTGAATTCCAGCAGGCCACGCTGCTGGCCTC | This paper |
| Luciferase + BCAP31 27 bp cloning PCR R | ACCAGTTCTTGCAAGCAGAATTCATTGGAGGCCAGCAGCGTGGCCT | This paper |
| Luciferase + SLC7A5 27 bp cloning PCR F | GGCAAGATCGCCGTGAATTCGGAGGCAGCCAAGCTCGTGGCCTG | This paper |
| Luciferase + SLC7A5 27 bp cloning PCR R | ACCAGTTCTTGCAAGCAGAATTCAGGCAGGCCACGAGCTTGGCTGC | This paper |
| Luciferase + STOML2 15 bp cloning PCR F | Same as Luciferase + STOML2 99 bp cloning PCR1 F | This paper |
| Luciferase + STOML2 15 bp cloning PCR R | CAGTTCTTGCAAGCAGAATTCAGTCACAGCCGACTGGAATTCACGGCGATCTTGCC | This paper |
| Luciferase + YKT6 15 bp cloning PCR F | Same as Luciferase + STOML2 99 bp cloning PCR1 F | This paper |
| Luciferase + YKT6 15 bp cloning PCR R | CAGTTCTTGCAAGCAGAATTCCACTCCCAGCTGACCGAATTCACGGCGATCTTGCC | This paper |

| | | |
|---|---|---|
| Luciferase + BCAP31 15 bp cloning PCR F | Same as Luciferase + STOML2 99 bp cloning PCR1 F | This paper |
| Luciferase + BCAP31 15 bp cloning PCR R | ACCAGTTCTTGCAAGCAGAATTCGGCCAGCAGCGTGGCGAATTCACGGCGATCTTGCC | This paper |
| Luciferase + SLC7A5 15 bp cloning PCR F | Same as Luciferase + STOML2 99 bp cloning PCR1 F | This paper |
| Luciferase + SLC7A5 15 bp cloning PCR R | ACCAGTTCTTGCAAGCAGAATTCGCCACGAGCTTGGCTGAATTCACGGCGATCTTGCC | This paper |
| Luciferase + GCTG cloning PCR F | GAAGGGCGGCAAGATCGCCGTGAATTCGCTGGAATTCTGCTTG | This paper |
| Luciferase + GCTG cloning PCR R | GAACCAGTTCTTGCAAGCAGAATTCCAGCGAATTCACGGCGATC | This paper |
| Luciferase + STOML2 15 bp GC-unpaired PCR F | Same as Luciferase + STOML2 99 bp cloning PCR1 F | This paper |
| Luciferase + STOML2 15 bp GC-unpaired PCR R | CAGTTCTTGCAAGCAGAATTGACTGACAGCCGACTGGAATTCACGGCGATCTTGCC | This paper |
| Luciferase + STOML2 15 bp GC-repaired PCR F | Same as Luciferase + STOML2 99 bp cloning PCR1 F | This paper |
| Luciferase + STOML2 15 bp GC-repaired PCR R | CAGTTCTTGCAAGCAGAATTGACTGACAGCCCAGTCGAATTCACGGCGATCTTGCC | This paper |
| Luciferase + YKT6 51 bp GC-unpaired PCR1 F | Same as Luciferase + STOML2 99 bp cloning PCR1 F | This paper |
| Luciferase + YKT6 51 bp GC-unpaired PCR1 R | CCAGCTCACCATGGAAAGACGAGGAAAAGAATTCACGGCGAT | This paper |
| Luciferase + YKT6 51 bp GC-unpaired PCR2 F | CGTCTTTCCATGGTGAGCTGGGAGTGTCCATCATC | This paper |
| Luciferase + YKT6 51 bp GC-unpaired PCR2 R | CCAGTTCTTGCAAGCAGAATTC | This paper |
| Luciferase + YKT6 51 bp GC-repaired PCR F | Same as Luciferase + STOML2 99 bp cloning PCR1 F | This paper |
| Luciferase + YKT6 51 bp GC-repaired PCR R | GTTCTTGCAAGCAGAATTCTCTTCCCCTCATCATCCACACTGCCAGCTCACCATGGAAAG | This paper |
| Luciferase + STOML2 intron cloning PCR1 F | Same as Luciferase + STOML2 99 bp cloning PCR1 F | This paper |
| Luciferase + STOML2 intron cloning PCR1 R | GTCAGTAAGACCAATAGGTGC | This paper |
| Luciferase + STOML2 intron cloning PCR2 F | GCACCTATTGGTCTTACTGACATATGTGCAGAGTCTCAAGG | This paper |
| Luciferase + STOML2 intron cloning PCR2 R | GTGGAGAGAAAGGCAAAGTGGACAGGTAAAGGACTCCATCGAT | This paper |
| Luciferase + STOML2 intron cloning PCR3 F | TCCACTTTGCCTTTCTCTCCAC | This paper |
| Luciferase + STOML2 intron cloning PCR3 R | CTGAACCAGTTCTTGCAAGCAGAATTCACGGCGATCTTGCCGCCCTTCTTG | This paper |
| Luciferase + YKT6 intron cloning PCR1 F | Same as Luciferase + STOML2 99 bp cloning PCR1 F | This paper |
| Luciferase + YKT6 intron cloning PCR1 R | Same as Luciferase + STOML2 intron cloning PCR1 R | This paper |
| Luciferase + YKT6 intron cloning PCR2 F | GCACCTATTGGTCTTACTGACAAACTGAGCCCAGGAAAGGGGGA | This paper |
| Luciferase + YKT6 intron cloning PCR2 R | GTGGAGAGAAAGGCAAAGTGGAGCAGCTGAGTGCACCTGTGTTT | This paper |
| Luciferase + YKT6 intron cloning PCR3 F | Same as Luciferase + STOML2 intron cloning PCR3 F | This paper |
| Luciferase + YKT6 intron cloning PCR3 R | Same as Luciferase + STOML2 intron cloning PCR3 R | This paper |
| pSJ560-TN/CA/7-PA(ΔX) cloning PCR1 F | GGAGACCCAAGCTGTTAACGCTAGCAGTTAACCGGAGTAC | This paper |
| pSJ560-TN/CA/7-PA(ΔX) cloning PCR1 R | CTCTTCGCCTCTTTCGGACTGACGGAAGGAATCCCATAGACTCCTACT | This paper |

| | | |
|---|---|---|
| pSJ560-TN/CA/7-PA(ΔX) cloning PCR2 F | CAGTCCGAAAGAGGCGAAGAGACAATAGAAGAAAAATTTGAGATTACAGG | This paper |
| pSJ560-TN/CA/7-PA(ΔX) cloning PCR2 R | AATGTCCTGTAGCTCTGCTAGCACCTGCTTCCAAGCCATGAG | This paper |
| pHW-Perth09-PA(ΔX) cloning PCR1 F | Same as pSJ560-H1N1pdm09-PA(ΔX) cloning PCR1 F | This paper |
| pHW-Perth09-PA(ΔX) cloning PCR1 R | CTTCTATTGTTTCTTCGCCTCTTTCGGACTGACGGAAGG | This paper |
| pHW-Perth09-PA(ΔX) cloning PCR2 F | CCTTCCGTCAGTCCGAAAGAGGCGAAGAAACAATAGAAG | This paper |
| pHW-Perth09-PA(ΔX) cloning PCR2 R | TCGCATCATATAATGGGATCCCTTCTCCTTCGTGACTTGG | This paper |
| pHW197-PR8-M-STOML2 cloning PCR1 F | ATGCCCTTAATGGGAACGGGGATC | This paper |
| pHW197-PR8-M-STOML2 cloning PCR1 R | TTACTCCAGCTCTATGCTGAC | This paper |
| pHW197-PR8-M-STOML2 cloning PCR2 F | GTCAGCATAGAGCTGGAGTAAGTCATCAACGTGCCTGAGCAG | This paper |
| pHW197-PR8-M-STOML2 cloning PCR2 R | GATATTTGCGGCAATAGTGAGACAGAGTTACATTGTCGAG | This paper |
| pHW197-PR8-M-STOML2 cloning PCR3 F | TCTCACTATTGCCGCAAATATCATTGGG | This paper |
| pHW197-PR8-M-STOML2 cloning PCR3 R | CATTTTGGGCCGCCGGGTTATTAGTAGAAACAAGGTAGTT | This paper |
| pHW197-PR8-M-STOML2 cloning PCR4 F | AATAACCCGGCGGCCCAAAATG | This paper |
| pHW197-PR8-M-STOML2 cloning PCR4 R | Same as pHW196-PR8-NA-STOML2 cloning PCR4 R | This paper |
| pHW197-PR8-M-STOML2-mut cloning F | Same as pHW196-PR8-NA-STOML2-mut cloning F | This paper |
| pHW197-PR8-M-STOML2-mut cloning R | Same as pHW196-PR8-NA-STOML2-mut cloning R | This paper |


1. Gaucherand, L. *et al.* The Influenza A Virus Endoribonuclease PA-X Usurps Host mRNA Processing Machinery to Limit Host Gene Expression. *Cell Reports* **27**, 776-792.e7 (2019).
2. Sievers, F. *et al.* Fast, scalable generation of high-quality protein multiple sequence alignments using Clustal Omega. *Molecular Systems Biology* **7**, 539 (2011).
3. Crooks, G. E., Hon, G., Chandonia, J. M. & Brenner, S. E. WebLogo: A Sequence Logo Generator. *Genome Research* **14**, 1188–1190 (2004).
4. Hu, T. *et al.* Variant G6PD levels promote tumor cell proliferation or apoptosis via the STAT3/5 pathway in the human melanoma xenograft mouse model. *BMC Cancer* **13**, 251 (2013).

5. Khaperskyy, D. A., Schmaling, S., Larkins-Ford, J., McCormick, C. & Gaglia, M. M. Selective Degradation of Host RNA Polymerase II Transcripts by Influenza A Virus PA-X Host Shutoff Protein. *PLoS Pathog* **12**, e1005427 (2016).
6. Slaine, P. D. *et al.* Thiopurines Activate an Antiviral Unfolded Protein Response That Blocks Influenza A Virus Glycoprotein Accumulation. *J Virol* **95**, e00453-21 (2021).

### Extended Data Fig. 1

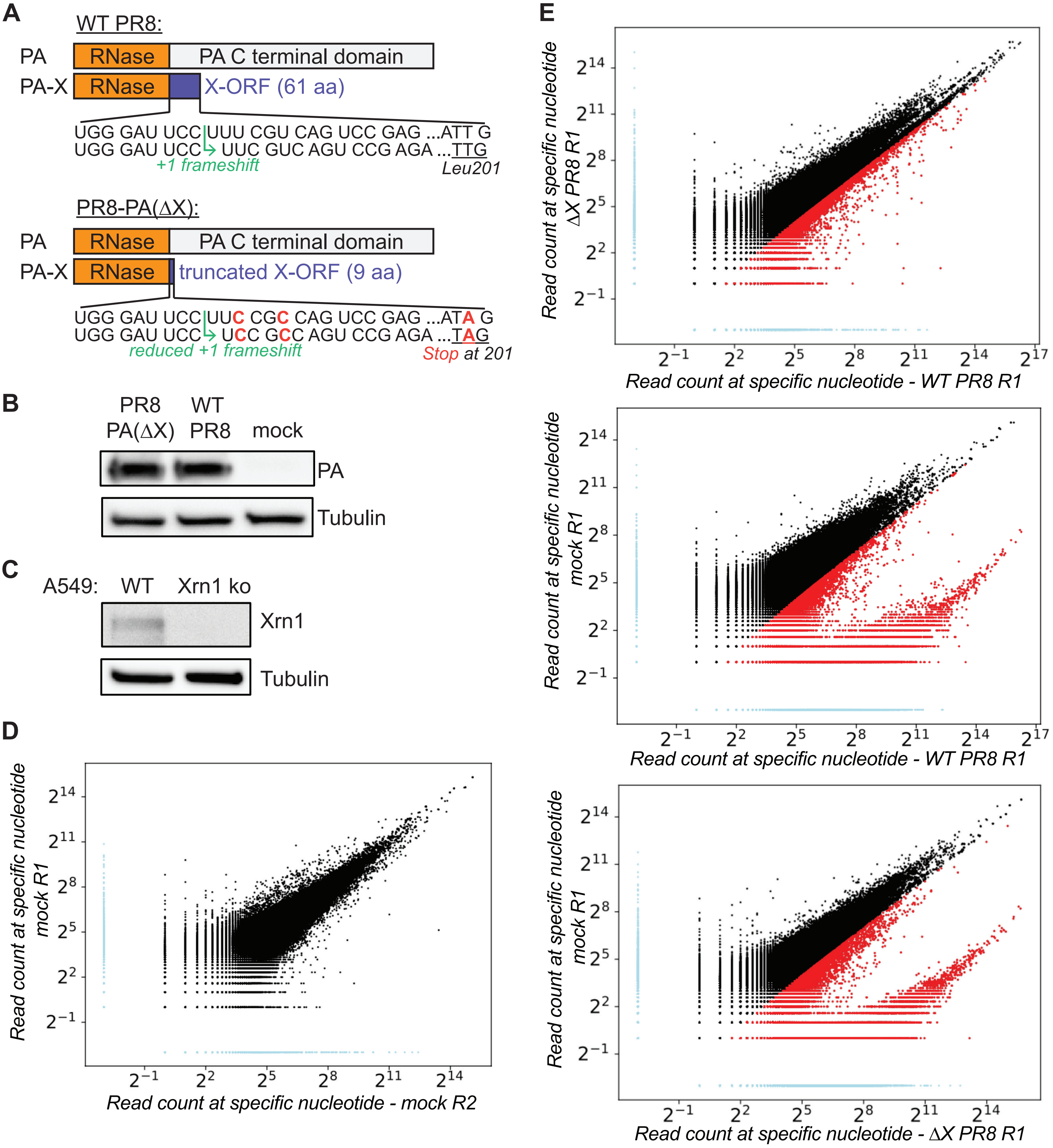

***Extended Data Fig. 1: Characteristics of the system used to identify PA-X cut sites transcriptome-wide.*** *(A) Strategy used to engineer a virus that lacks PA-X, PR8-PA(ΔX), compared to WT PR8. Adapted from*[1]*. (B) Protein lysates of Xrn1 knock out (ko) A549 cells infected with WT PR8 or PR8-PA(ΔX), or mock infected, were probed with antibodies against PR8 PA or β-tubulin as a loading control. Images are representative of 2 experiments. (C) Protein lysates of WT or Xrn1 knock out A549 cells were probed with antibodies against Xrn1 or β-tubulin as a loading control, to check for loss of Xrn1. Images are representative of 2 experiments. (D-E) Representation of individual chromosomal positions in the 5' RACE-seq data. For each sample, reads with their 5' end mapping to the same nucleotide were counted and plotted to compare different datasets: (D) mock samples from replicate 1 vs. replicate 2, (E) replicate 1 WT PR8 vs. PR8-PA(ΔX) (top), WT PR8 vs. mock infected (middle) and PR8-PA(ΔX) vs. mock infected (bottom). For each plot, light blue dots correspond to locations that are unique to one sample, while red and black dots correspond to locations that are common in the two samples. Red dots represent locations that have two-fold or more reads in the sample on the x axis* vs. *than in the sample on the y axis. Similar plots were obtained when comparing other replicates.*

**Extended Data Fig. 2**

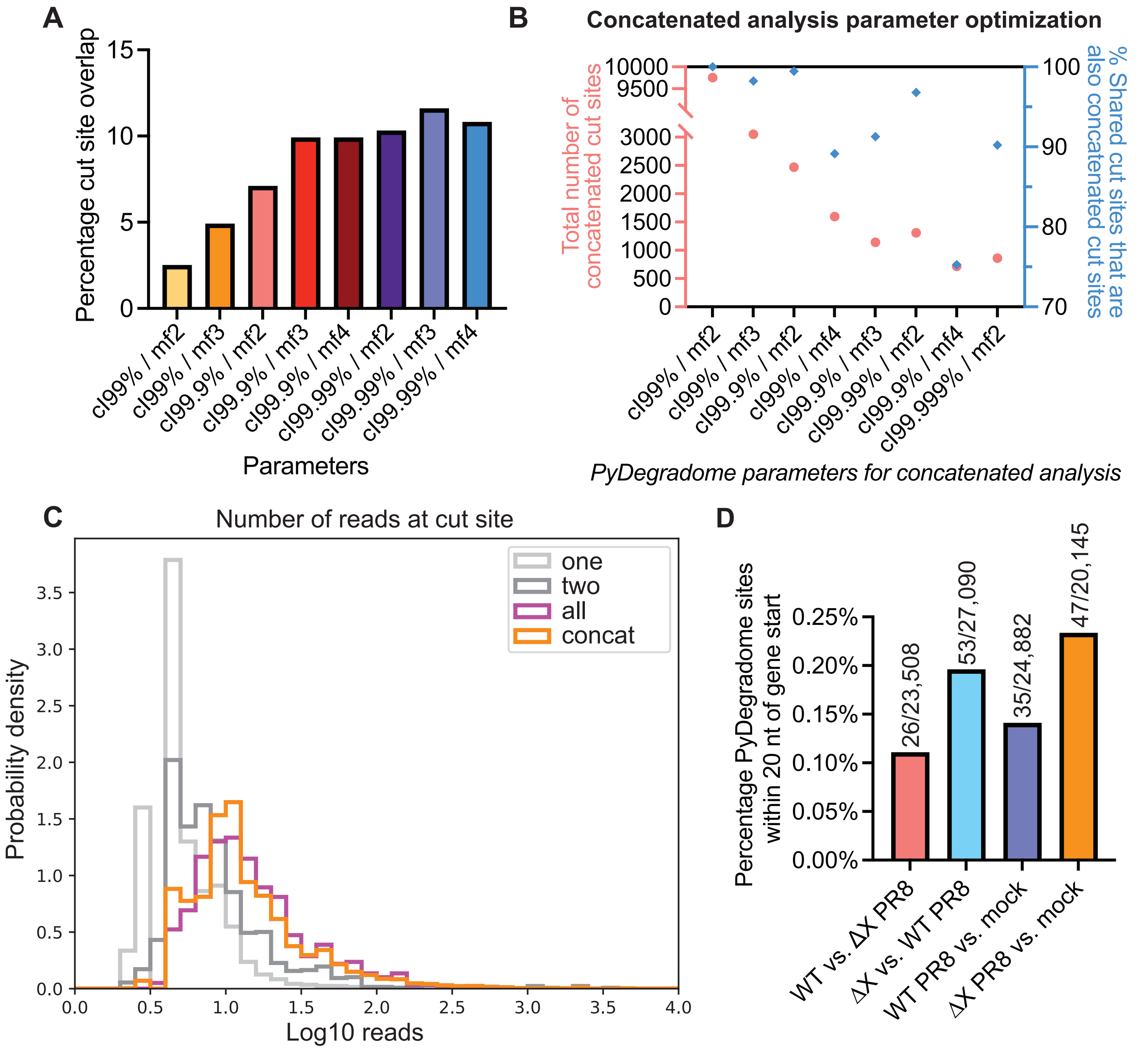

***Extended Data Fig. 2: Optimization of parameters for the identification of PA-X cut sites transcriptome-wide and analysis of output characteristics.*** *(A-B) PyDegradome was run for each replicate (A) or on concatenated samples (B) on WT PR8* vs. *PR8-PA(ΔX) infection samples using different parameters as indicated on the x axis (see Methods). cl = confidence levels, mf = multiplicative factor. The window of analysis was 4 nucleotides in all cases. In A, the percentage of cut sites shared between all three replicates was plotted for each set of parameters. In B, the total number of cut sites identified by PyDegradome (red circles, left y axis) and the percentage of shared cut sites also found as concatenated cut sites (blue diamonds, right y axis) were plotted for each set of parameters. (C) Histogram of the number of reads at the cut site for PA-X sites (PyDegradome comparison: WT PR8* vs. *PR8-PA(ΔX)) in one (light grey), two (dark grey), or all three replicates ("all", magenta), or through the concatenated analysis ("concat", orange). (D) Percentage of sites identified by PyDegradome in at least one replicate that are located within the first 20 nucleotides of a gene when comparing WT PR8* vs. *PR8-PA(ΔX), PR8-PA(ΔX)* vs. *WT PR8, WT PR8* vs. *mock and PR8-PA(ΔX)* vs. *mock.*

**Extended Data Fig. 3**

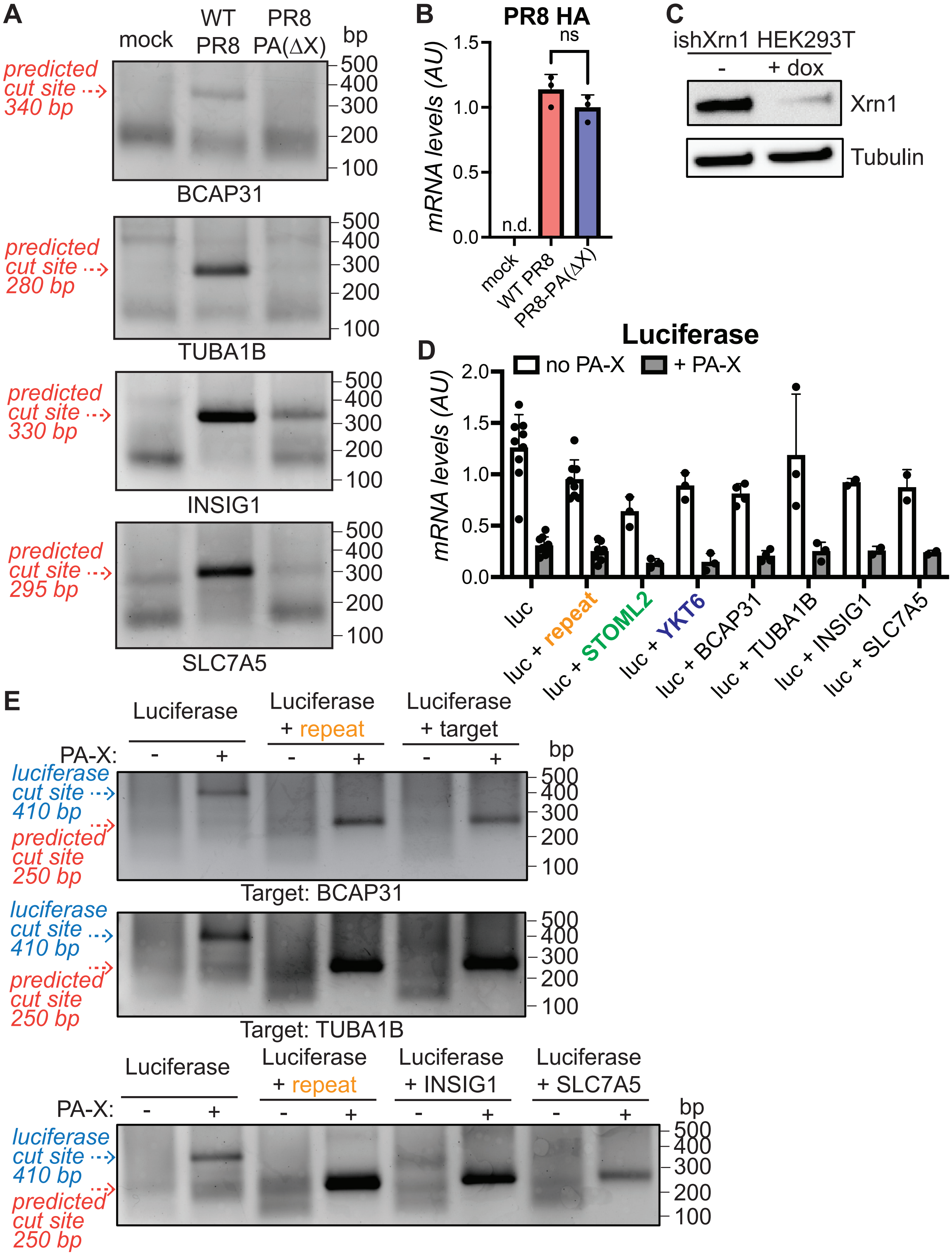

***Extended Data Fig. 3: Further validation of PA-X cut sites identified by PyDegradome.*** *(A-B) Xrn1 knock out A549 cells were infected with WT PR8 or PR8-PA(∆X), or mock infected. (A) RNA was isolated, 5' RACE was performed using primers specific for BCAP31, TUBA1B, INSIG1 or SLC7A5, and the PCR products were run on an agarose gel. Primers were positioned ~ 200-300 nucleotides downstream of the predicted cut sites. The predicted size of DNA bands coming from cut sites identified by PyDegradome are indicated by the red dotted arrows. (B) PR8 HA RNA levels were quantified by qRT-PCR, normalized to 18S and plotted as mean ± standard deviation. AU, arbitrary units; n.d. not defined; ns, not significant, One-way ANOVA with Dunnett's multiple comparison test. n = 3. (C) HEK293T ishXrn1 cells were treated with no drug or doxycycline for 3-4 days to induce shRNAs against Xrn1, then protein lysates were collected and probed with antibodies against Xrn1, or β-tubulin as a loading control, to check for efficient knock down of Xrn1. Images are representative of 3 experiments. (D-E) HEK293T ishXrn1 cells were treated with doxycycline for 3-4 days to induce knock down of Xrn1, then transfected with luciferase reporters containing 99 bp insertions from the indicated genes, and where indicated, with PR8 PA-X. (D) RNA was extracted and luciferase mRNA levels were quantified by qRT-PCR, normalized to 18S and plotted as mean ± standard deviation. AU, arbitrary units. n ≥ 2. (E) The RNA was also used to run 5' RACE. Expected sizes of DNA bands coming from cut sites in the introduced target sequences are indicated by the red arrows, while the blue arrow in D indicates the size of the original luciferase cut site fragment. For all gels, the DNA bands were purified and sequenced to confirm their identities, and images are representative of 3 experiments or 2 experiments for the luciferase + INSIG1 and + SLC7A5 reporters. The SLC7A5 sequence was only consistently cut when inside the luciferase reporter.*

**Extended Data Fig. 4**

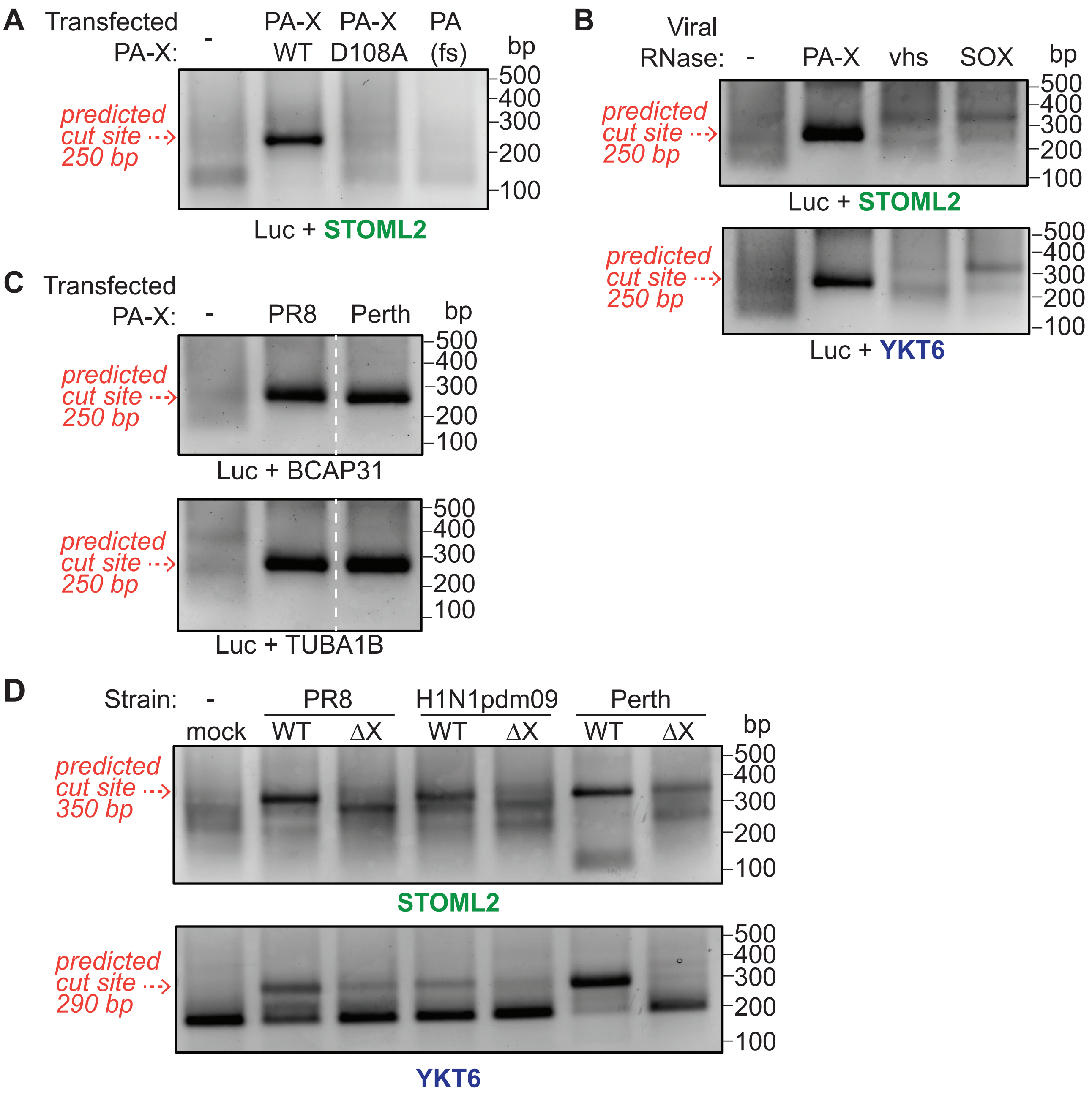


**E**

```
H1N1pdm09_PA-X:       MEDFVRQCFNPMIVELAEKAMKEYGEDPKIETNKFAAICTHLEVCFMYSDFHFIDERGES     60
PR8_PA-X              MEDFVRQCFNPMIVELAEKTMKEYGEDLKIETNKFAAICTHLEVCFMYSDFHFINEQGES     60
Perth_PA-X:           MEDFVRQCFNPMIVELAEKAMKEYGEDLKIETNKFAAICTHLEVCFMYSDFHFINEQGES     60
                      *******************:******* **************************:*:***

H1N1pdm09_PA-X:       IIVESGDPNALLKHRFEIIEGRDRIMAWTVVNSICNTTGVEKPKFLPDLYDYKENRFIEI     120
PR8_PA-X              IIVELGDPNALLKHRFEIIEGRDRTMAWTVVNSICNTTGAEKPKFLPDLYDYKENRFIEI     120
Perth_PA-X:           IVVELDDPNALLKHRFELIEGRDRTMAWTVVNSICNTTGAGKPKFLPDLYDYKENRFIEI     120
                      *:** .***********:****** **************. *******************

H1N1pdm09_PA-X:       GVTRREVHIYYLEKANKIKSEKTHIHIFSFTGEEMATKADYTLDEESRARIKTRLFTIRQ     180
PR8_PA-X              GVTRREVHIYYLEKANKIKSEKTHIHIFSFTGEEMATKADYTLDEESRARIKTRLFTIRQ     180
Perth_PA-X:           GVTRREVHIYYLEKANKIKSENTHIHIFSFTGEEMATKADYTLDEESRARIKTRLFTIRQ     180
                      *********************:**************************************

H1N1pdm09_PA-X:       EMASRSLWDSFVSPKEAKRQLKKNLRLQELCASLPTKVSHRTSPALKTLEPM--------     232
PR8_PA-X              EMASRGLWDSFVSPREEKRQLKKGLKSQEQCVSLPTKVSRRTSPALKILEPMWMDSNRTA     240
Perth_PA-X:           EMANRGLWDSFVSPKEAKKQLKKNLKSQELCVGLPTKVSHRTSPALRILEPMWMDSNRTA     240
                      ***.*.********:* *:****.*: ** *..******:******: ****

H1N1pdm09_PA-X:       ------------        232
PR8_PA-X              TLRASCLKCPKK        252
Perth_PA-X:           ALRASFLKCPKK        252
```

***Extended Data Fig. 4: The cut sites identified by PyDegradome are specific to PA-X and conserved across multiple influenza strains.*** *(A-C) HEK293T ishXrn1 cells were treated with doxycycline for 3-4 days to induce knock down of Xrn1, then transfected with luciferase reporters containing 99 bp insertions from the indicated genes, and where indicated, with WT PA-X from the PR8 strain (A-C), the catalytic mutant PR8 PA-X D108A (A), PR8 PA with a mutation to reduce frameshifting and prevent PA-X production (PA(fs)) (A), herpes simplex virus 1 (HSV-1) vhs or Kaposi's sarcoma-associated herpes virus (KSHV) SOX (B), or WT PA-X from the Perth influenza strain (C). RNA was extracted and used to run 5' RACE. Expected sizes of DNA bands coming from cut sites in the introduced target sequences are indicated by the red dotted arrows. (D) Xrn1 knock out A549 cells were infected with WT PR8 or PR8-PA(ΔX), WT H1N1pdm09 or H1N1pdm09-PA(ΔX), WT Perth or Perth-PA(ΔX), or mock infected. 5' RACE was then performed using primers specific for STOML2 or YKT6 ~250-300 nt downstream of the predicted cut sites. The PCR products were run on an agarose gel. The predicted size of DNA bands coming from cut sites identified by PyDegradome are indicated by the red dotted arrows. For all gels, the DNA bands were purified and sequenced to confirm their identities, and images are representative of 3 experiments. (E) Protein alignment of PA-X from the three different influenza strains PR8, H1N1pdm09 and Perth, generated using Clustal Omega[2].*

**Extended Data Fig. 5**

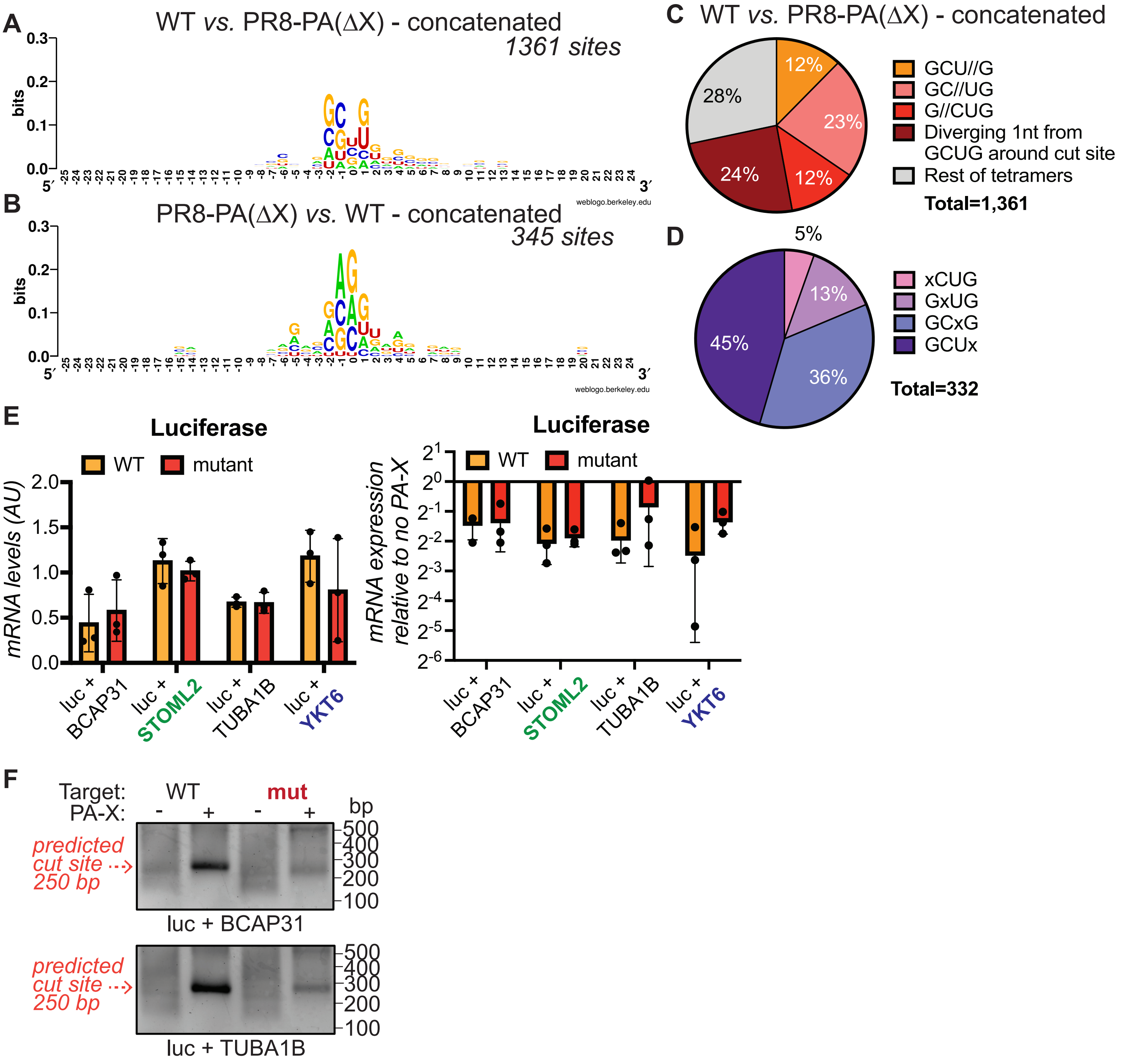

***Extended Data Fig. 5: PA-X preferentially cleaves RNA at GCUG tetramers based on the concatenated cut site analysis.*** *(A-B) WebLogo[3] representation of base enrichment around PA-X cut sites (WT PR8 vs. PR8-PA(∆X), A) or around control sites enriched in the PR8-PA(∆X) sample (PR8-PA(∆X) vs. WT PR8, B) for sites predicted by PyDegradome using the concatenated approach. (C) Percentage of PA-X cut sites containing GCUG or a tetramer with one nucleotide difference from GCUG, for sites identified by PyDegradome using the concatenated approach. // indicates the location of the cut, i.e. GCU//G indicates that PA-X cuts between the U and the G. (D) Further breakdown of the PA-X cut sites containing a tetramer with one nucleotide difference from GCUG around the cut site (marked by an x). (E-F) HEK293T ishXrn1cells were treated with doxycycline for 3-4 days to induce knock down of Xrn1, then transfected with luciferase reporters containing 99 bp insertions from the indicated genes, with or without PR8 PA-X. (E) RNA was extracted and luciferase mRNA levels were quantified by qRT-PCR, normalized to 18S and plotted as mean ± standard deviation. Left plot shows mRNA levels of each reporter in the absence of PA-X. Right plot shows mRNA levels of each reporter in the presence vs. absence of PA-X. AU, arbitrary units. n = 3. (F) The RNA was also used to run 5' RACE. Expected sizes of DNA bands coming from cut sites in the introduced target sequences are indicated by the red dotted arrows. For both gels, DNA bands were purified and sequenced to confirm their identities, and images are representative of 3 experiments*

### Extended Data Fig. 6

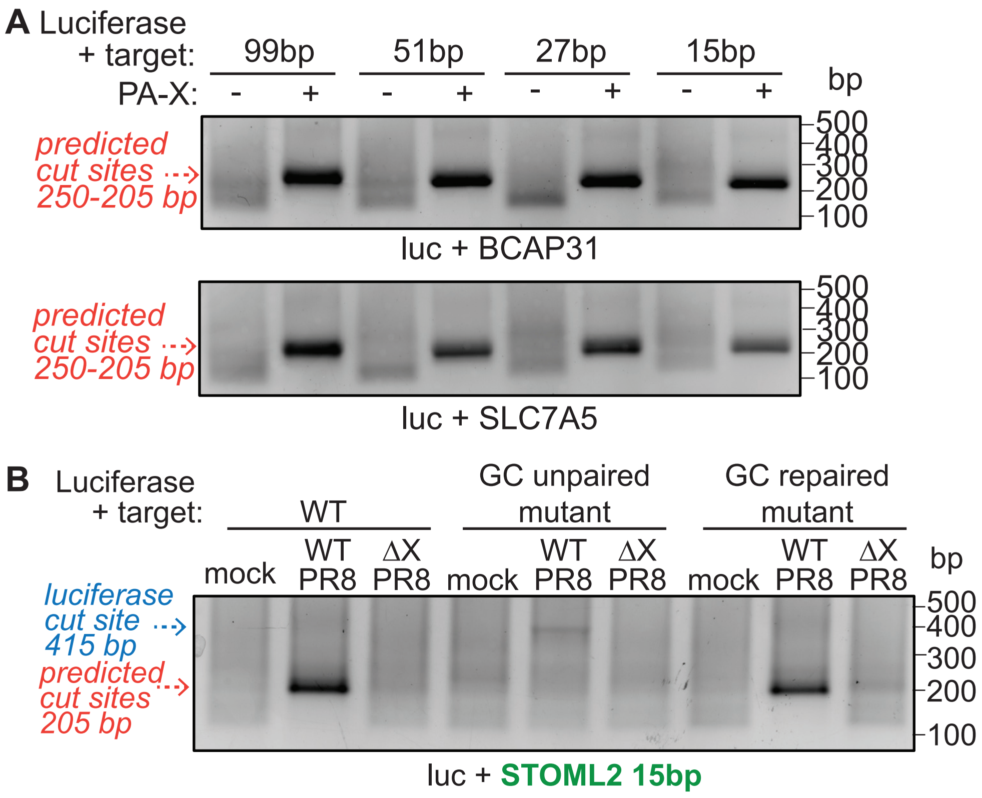


***Extended Data Fig. 6: PA-X preferentially cleaves RNA within a hairpin loop structure in transfected and infected cells.*** *(A) HEK293T ishXrn1cells were treated with doxycycline for 3-4 days to induce knock down of Xrn1, then transfected with luciferase reporters containing insertions of the indicated lengths from the BCAP31 and SLC7A5 genes, with or without PR8 PA-X. RNA was extracted and used to run 5' RACE. Expected sizes of DNA bands coming from cut sites in the introduced target sequences are indicated by the red dotted arrows (~250 bp for 99 bp constructs, ~220 bp for 51 bp constructs, ~210 bp for 27 bp constructs and ~205 bp for the 15 bp constructs). (B) HEK293T ishXrn1cells were treated with doxycycline for 3-4 days to induce knock down of Xrn1, then transfected with luciferase reporters containing 15 bp insertions from the STOML2 gene with or without the indicated mutations. 24 hours post transfection, cells were infected with WT PR8 or PR8-PA(ΔX), or mock infected overnight. RNA was then extracted and used to run 5' RACE. Expected sizes of DNA bands coming from cut sites in the introduced target sequences are indicated by the red dotted arrow, while the blue arrow indicates the size of the original luciferase cut site fragment. For all gels, the DNA bands were purified and sequenced to confirm their identities, and images are representative of 3 experiments (B) or 2 experiments (A).*

**Extended Data Fig. 7**

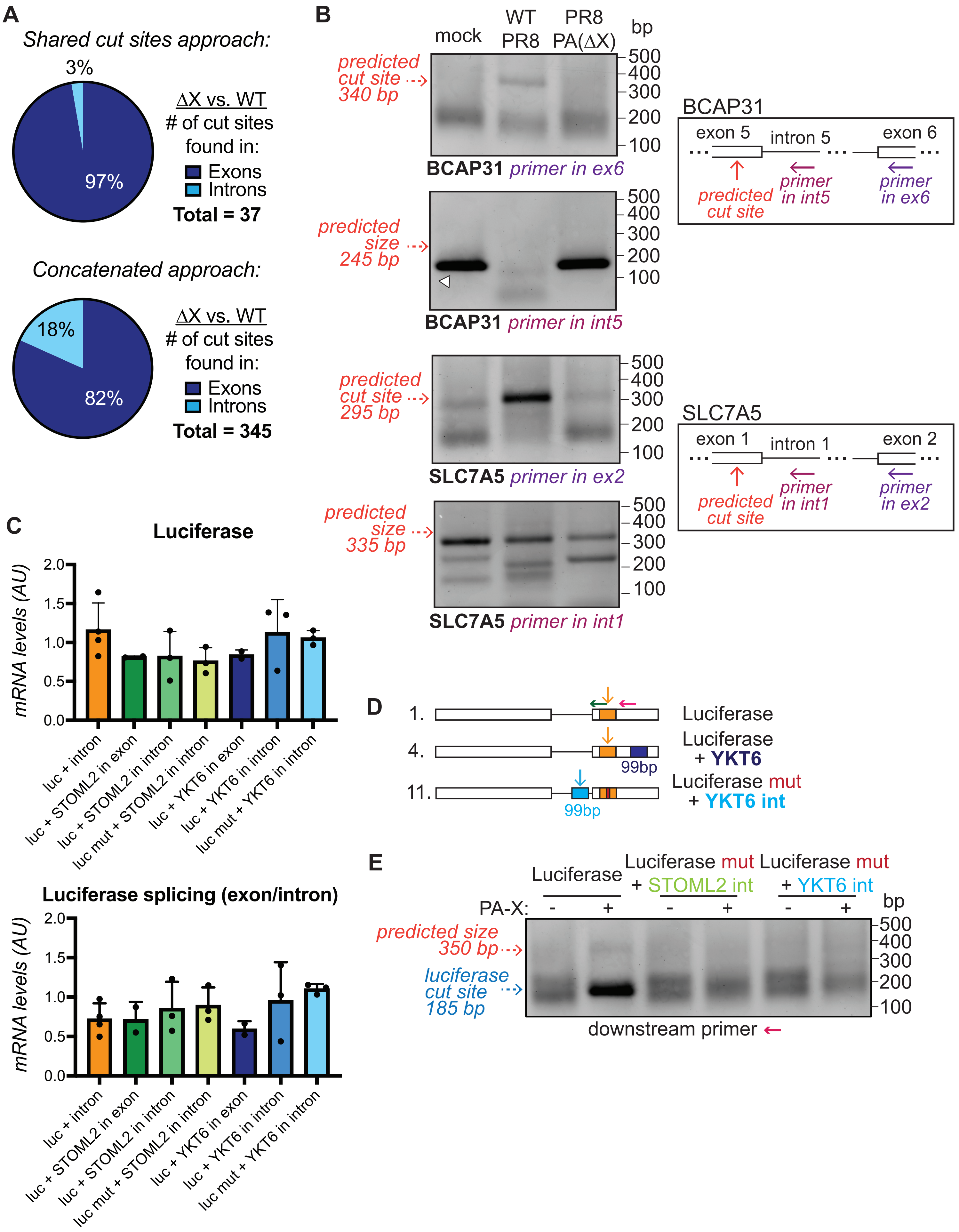

***Extended Data Fig. 7: PA-X preferentially cleaves RNAs within exons.*** *(A) Percentage of PR8-PA(ΔX) specific fragments found within introns or exons, for sites identified by PyDegradome using the shared cut sites approach (top) or the concatenated approach (bottom). (B) Xrn1 knock out A549 cells were infected with WT PR8 or PR8-PA(ΔX), or mock infected, and 5' RACE was performed on the RNA. Boxes show diagrams of the BCAP31 and SLC7A5 genes and positions of the reverse primers for 5' RACE. PCR products were run on an agarose gel. Top gels using reverse primers in exons (dark purple primers) are the same gels as* ***Extended Data Fig. 3A*** *and are included for comparison. Red dotted arrows indicate the size of fragments originating from previously validated cut sites. Bottom gels show products obtained using reverse primers in the intron (light purple primers), and red dotted arrows indicate the predicted size of PCR products that would appear if PA-X cleaved unspliced pre-mRNAs. White arrowhead indicates a fragment that map to an exon/intron junction. New gel images are representative of 3 experiments. (C-E) Testing of luciferase reporters with sequence insertions in the introns, as shown in diagram in D. HEK293T ishXrn1 cells were treated with doxycycline for 3-4 days to induce knock down of Xrn1, then transfected with luciferase reporters containing the 99 bp insertions in the indicated genes either in an exon or an intron, with or without PR8 PA-X. (C) RNA was extracted and luciferase mRNA levels were quantified by qRT-PCR, normalized to 18S and plotted as mean ± standard deviation. AU, arbitrary units; n ≥ 2. The top graph shows levels of the processed mRNA (obtained using two primers that bind in exons). The bottom graph shows levels of the unspliced pre-mRNA (obtained using one primer in an exon and one primer in the intron). (D-E) RNA was also used to run 5' RACE. (D) Green and magenta arrows indicate positions of 5' RACE PCR reverse primers, vertical arrows indicate location of predicted cut sites. (E) PCR products were obtained using the magenta arrow primer from (D), then were separated on an agarose gel. Blue dotted arrow indicates size of PCR products originating from the original luciferase cut site, red dotted arrow indicates the predicted size of PCR products that would originate from the sequence inserted in the introns. Gel images are representative of 3 experiments.*

**Extended Data Fig. 8**

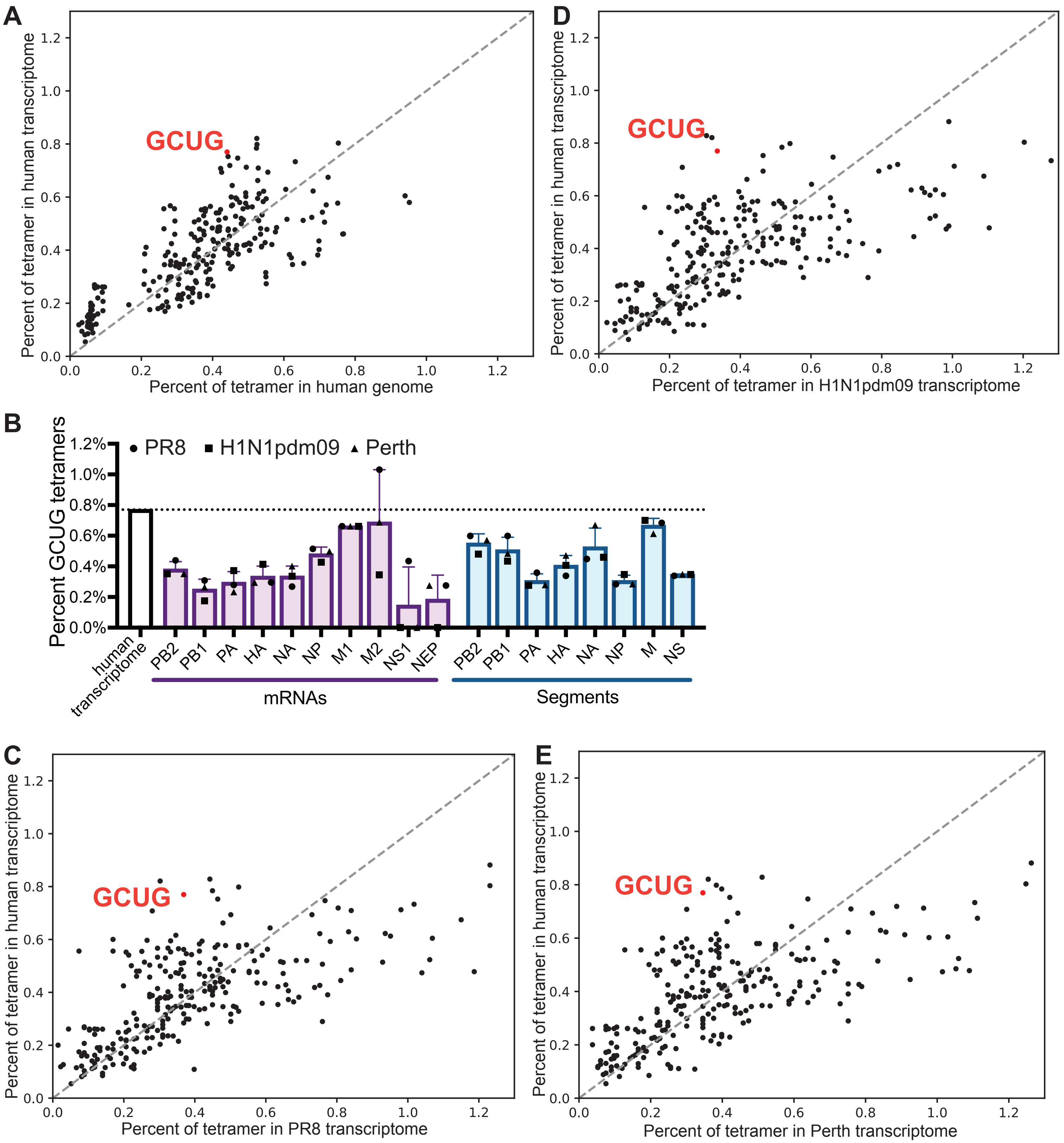

***Extended Data Fig. 8: GCUG tetramers are enriched in the human transcriptome.*** *The percentage of tetramers in the human and viral genomes and transcriptomes was calculated by counting the number of each tetramer and dividing it by the total number of tetramers in the sequence (i.e. length of the sequence minus 3). Percentages for each tetramer are plotted to visualize which tetramers are more abundant in the human genome vs. transcriptome (A) and the human transcriptome vs. the transcriptome of 3 influenza A virus strains (C-E). Each dot represents a specific tetramer, with the red dot representing the GCUG tetramer. (B) The percentage of GCUG tetramers is plotted for each influenza mRNA (i.e. the positive strand, purple), and for each influenza genomic RNA (i.e. negative strand, blue). Each symbol represents a different influenza strain.*

**Extended Data Fig. 9**

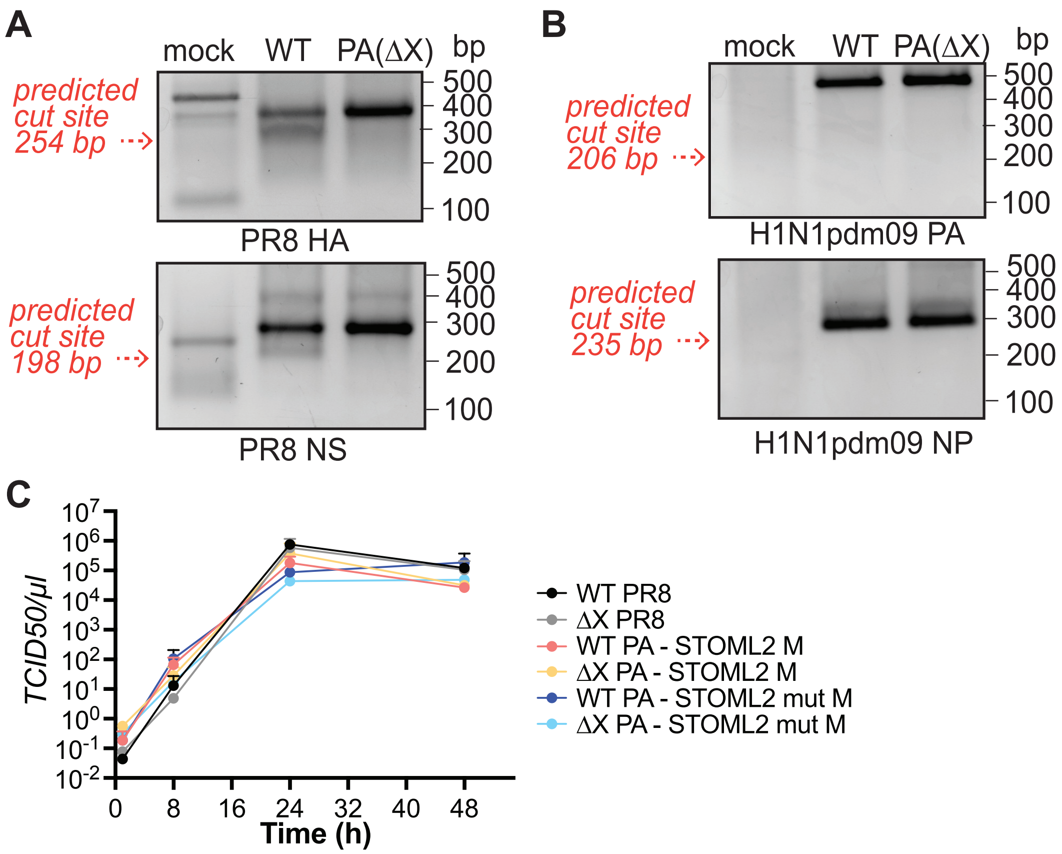


***Extended Data Fig. 9: Viral mRNAs are not efficiently cleaved by PA-X.*** *(A-B) Xrn1 knock out A549 cells were infected with WT PR8 or PR8-PA(ΔX) (A), or WT H1N1pdm09 or H1N1pdm09-PA(ΔX) (B), or mock infected. RNA was extracted to run 5' RACE using primers ~150-200 nt downstream of GCUG sites that are predicted to be inside a hairpin loop within the indicated viral mRNAs. Red dotted arrows indicate the predicted sizes of PCR products that would originate from cleavage at these GCUG sites. Gel images are representative of 3 experiments. (C) MDCK cells were infected at MOI 0.05 with the indicated viruses (see* ***Fig. 6F*** *for details on virus construction). Supernatants were collected at 1, 8, 24 and 48 hours post infection and viral titers were quantified by TCID50. n = 2.*
